# Restoration of deficient IL-15 signaling enhances innate lymphoid cell immunity in brain metastasis

**DOI:** 10.64898/2026.10.02.756334

**Authors:** Timothy P. McMullen, Pascal Naef, Erika Zagni, Lei Tian, Eva Zhao, Noah Wechter, Aaron J. Longworth, Isam Adam, Jacob Insua-Rodriguez, Hannah Savage, Isaac Hur, Sophia Aziza Garcia, Dennis Ma, Michael H. Liew, Sharmila Mallya, Kai Kessenbrock, Timothy E. O’Sullivan, Ahmed Mohyeldin, Jianhua Yu, Devon A. Lawson

**Affiliations:** Department of Physiology and Biophysics, School of Medicine, University of California, Irvine; Medical Scientist Training Program, School of Medicine, University of California, Irvine; Sue and Bill Gross Stem Cell Research Center, School of Medicine, University of California, Irvine; Department of Biological Chemistry, School of Medicine, University of California, Irvine; Center for Complex Biological Systems, University of California, Irvine; Department of Microbiology, Immunology, and Molecular Genetics, University of California, Los Angeles; Division of Hematology & Oncology, Department of Medicine, School of Medicine, University of California, Irvine; Chao Family Comprehensive Cancer Center, University of California, Irvine

## Abstract

Natural killer (NK) cells and type 1 innate lymphoid cells (ILC1s), collectively known as group 1 innate lymphoid cells (ILCs), are critical mediators of anti-tumor immunity, yet their role in brain metastasis remains poorly defined. Using single-cell RNA sequencing and functional depletion models, we show that group 1 ILCs fail to provide adequate tumor control in breast cancer brain metastasis (BCBM). In contrast to the liver, where diverse, functionally competent group 1 ILC populations facilitate excellent metastatic control, group 1 ILCs in BCBM show limited diversity and profound tissue-specific dysfunction such that their depletion had no effect on tumor burden despite modest parenchymal infiltration. We attribute this group 1 ILC dysfunction to severely restricted IL-15 availability, with IL-15Rα protein detected at 500-fold lower concentration in the brain than in the liver. Local IL-15 complex administration restored group 1 ILC accumulation, cytotoxicity, and tissue residency, significantly reducing brain metastatic burden independent of T cells. Intracranial delivery of IL-15-secreting induced pluripotent stem-cell-derived (iPSC)-NK cells, already in clinical testing for non-small cell lung cancer, significantly reduced tumor burden. Collectively, these findings reveal how tissue-specific differences in cytokine availability shape group 1 ILC population diversity and disease response, establishing localized IL-15 signaling as a compelling strategy to unlock innate immunity against intracranial malignancies.

**One Sentence Summary:** Group 1 ILCs, including NK cells and ILC1s, can be pharmacologically enhanced with IL-15 to improve control of breast cancer brain metastasis.

## INTRODUCTION

Brain metastasis is a devastating complication of advanced breast cancer associated with substantial neurological morbidity and poor clinical outcomes^1^. The incidence of breast cancer brain metastasis (BCBM) is particularly high in HER2-positive and triple-negative breast cancer, affecting approximately one-third of patients with metastatic disease in these subtypes^2^. Current treatment strategies include surgical resection, stereotactic radiosurgery, whole-brain radiotherapy, and increasingly, central nervous system (CNS)-active systemic therapies^3–5^. Although these approaches can provide temporary disease control, intracranial recurrence remains common, and CNS-directed treatments can cause substantial neurological and cognitive toxicity^3,4,6^. Thus, there remains a critical need for therapeutic strategies that provide durable control of BCBM while limiting treatment-associated toxicity.

Besides targeting T-cells, natural killer (NK) cells are particularly attractive candidates for cancer immunotherapy because they recognize malignant cells without requiring prior antigen exposure or HLA-restricted antigen presentation^7–10^. This enables the development of allogeneic, “off-the-shelf” NK-cell therapies with a low risk of graft-versus-host disease and, in early clinical studies, limited cytokine release syndrome and neurotoxicity^11–13^. However, NK-cell therapies remain challenging in solid tumors because of limited tumor infiltration, persistence, and suppression within the tumor microenvironment^11,14,15^.

NK cells and type 1 innate lymphoid cells (ILC1s) comprise group 1 innate lymphoid cells (ILCs; **Fig. 1A**)^16,17^. NK cells are predominantly circulating and highly cytotoxic, whereas ILC1s are generally considered tissue-resident. In mice, NK cells are commonly characterized by CD49b and Eomes expression, whereas ILC1s preferentially express CD49a and depend on the transcriptional regulator Hobit, although substantial phenotypic heterogeneity exists across tissues and disease states^17–19^. In addition, CD49a^+^CD49b^+^ ILC1-like populations, also known as tissue-resident NK cells, can emerge in tumors in response to microenvironmental signals, including TGF-β, highlighting the considerable plasticity within the group 1 ILC compartment^20–23^.

**Figure 1.**
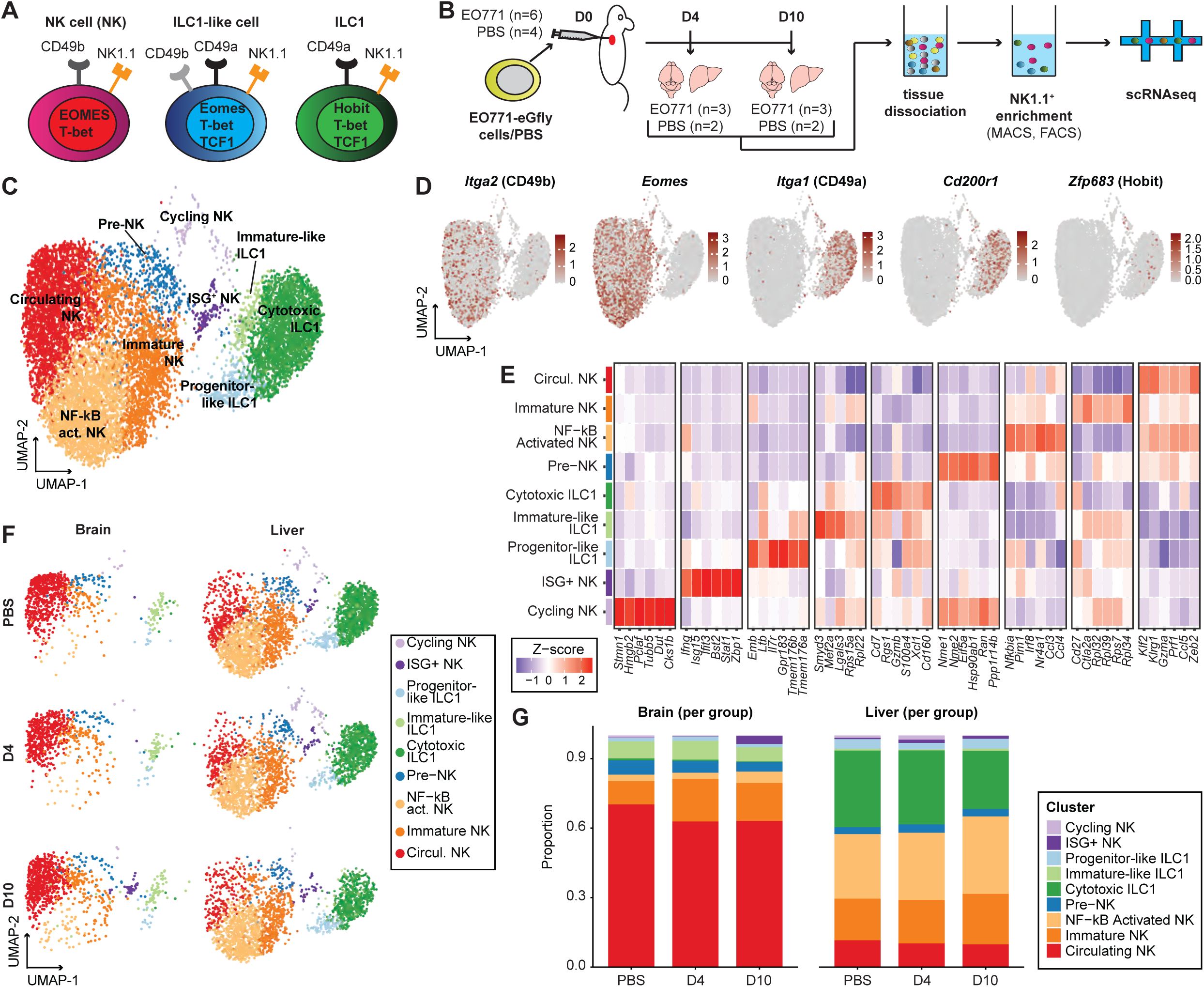
Group 1 ILC diversity is restricted in the CNS. **(A)** Schematic of surface receptors and transcription factors used to define group 1 innate lymphoid cells (ILCs), including NK cells, ILC1-like cells, and group 1 ILCs (ILC1s). **(B)** Experimental workflow for isolation and sequencing of metastasis-infiltrating NK1.1^+^ group 1 ILCs from brain and liver. Metastatic BL/6 mice (n=6) received EO771 eGFP/luciferase (eGfly) cells via intracardiac injection; control mice (n=4) received PBS. Mice were sacrificed at day 4 (D4; metastatic n=3, control n=2) and day 10 (D10; metastatic n=3, control n=2) post-injection. Brains and livers were enzymatically digested, and NK1.1^+^ cells were enriched by magnetic-activated cell separation (MACS) followed by fluorescence-activated cell sorting (FACS) prior to droplet-based scRNA-seq. **(C)** UMAP visualization of scRNA-seq profiles from sorted NK1.1^+^ cells isolated from the brain and liver of PBS-injected (healthy) or tumor-burdened BL/6 mice. **(D)** Feature plots showing the relative expression of *Itga2* (CD49b), *Eomes*, *Itga1* (CD49a), *Cd200r1,*and *Zfp683* (Hobit). **(E)** Heat map of Z-scored expression of NK-associated genes, ILC1-associated genes, cytotoxicity genes, and cell-cycle genes across clusters. **(F)** UMAP from (C) split by tissue (liver vs brain) and condition (PBS control vs tumor-burdened). **(G)** Bar plots showing the cluster distribution in brains (left panel) and livers (right panel) per group, either injected with PBS (Control group) or 4 and 10 days post intracardiac injection of EO771 cells (D4, D10).

NK-cell activity is regulated by integrating signals from germline-encoded activating and inhibitory receptors^24^. Healthy cells expressing MHC class I engage inhibitory NK-cell receptors, whereas malignant transformation can result in loss of MHC-I-mediated inhibition and increased expression of stress-induced ligands for activating receptors^25^. Cytokines including IL-2, IL-12, IL-15, IL-18, and type I interferons further regulate NK-cell survival, proliferation, priming, and effector function, with IL-15 playing a particularly important role in NK-cell development and homeostasis^26,27^. NK cells can additionally recognize antibody-coated tumor cells through Fc receptors and mediate antibody-dependent cellular cytotoxicity^28^. Once activated, NK cells directly eliminate malignant cells through perforin- and granzyme-mediated cytotoxicity and death-receptor pathways, including FasL and TRAIL, while secreting cytokines and chemokines such as IFN-γ, TNF, CCL3, CCL4, and CCL5 that shape broader antitumor immune responses^29–31^. ILC1s similarly produce IFN-γ and TNF and can acquire cytotoxic effector programs, including granzyme C expression in distinct murine ILC1 subsets^20,32,33^. However, group 1 ILC phenotype and function depend heavily on the tissue microenvironment, and tumor-associated ILC1-like states can show impaired antitumor activity^20–22,34^. Thus, although group 1 ILCs regulate cancer progression and NK cells can restrict metastatic spread at multiple stages of the metastatic cascade, how these populations are regulated and function in BCBM remains poorly understood^20,35^.

The brain represents a highly specialized immune environment. Although historically considered immune-privileged, the CNS is now recognized to undergo active but tightly regulated immune surveillance^36,37^. The blood-brain barrier (BBB) and other CNS interfaces regulate immune-cell trafficking, while resident neural, glial, and myeloid cells shape local immune responses^36–38^. Consequently, immune-cell recruitment and function within the brain differ substantially from peripheral tissues. NK cells can enter the CNS during inflammation and malignancy, but they represent a relatively small and poorly characterized component of the brain immune compartment^39,40^. How the brain microenvironment regulates group 1 ILC recruitment, phenotype, and antitumor activity, and whether these cells can be therapeutically harnessed to control BCBM, remains unclear.

Here, we use an immunocompetent mouse model of breast cancer metastasis to show that brain group 1 ILCs display markedly reduced cellular diversity compared with their counterparts in the liver, with a predominance of circulating NK-cell states and a relative paucity of cytotoxic ILC1 populations. Furthermore, depletion of NK1.1^+^ cells increases metastatic burden in the liver but has a negligible effect on metastatic burden in the brain, indicating a site-specific failure of group 1 ILC-mediated tumor surveillance in the brain. We identify limited IL-15 signaling as a potential constraint on group 1 ILC activity within the brain and show that intracranial administration of an IL-15/IL-15Rα complex expands and activates NK cells, effectively restoring control of brain metastasis. Finally, we show that intracranial delivery of human induced pluripotent stem cell (iPSC)-derived NK cells, which are genetically modified to secrete IL-15, similarly suppresses metastatic outgrowth in a xenograft model. Together, these findings identify the brain microenvironment as a critical determinant of group 1 ILC-mediated metastatic control and demonstrate that enhancing NK-cell activity can overcome this tissue-specific limitation.

## RESULTS

### Group 1 ILC diversity is restricted in the CNS

We investigated the group 1 ILC response to BCBM using single-cell RNA-sequencing (scRNA-seq). We used an experimental metastasis model in which EO771 murine breast cancer cells are injected into the left ventricle of C57BL/6J (BL/6) mice and disseminate via the arterial circulation to diverse tissues. We directly compared the response in the brain with that in the liver, where group 1 ILCs robustly restrict metastatic growth^41,42^. Single cells were isolated from healthy brains and livers (n = 4 BL/6 mice) or from metastasis-bearing brains and livers at 4 and 10 days after tumor inoculation (n = 3 BL/6 mice per time point). These cell isolates were enriched for group 1 ILCs by lineage depletion using magnetic-activated cell sorting (MACS), followed by fluorescence-activated cell sorting (FACS) of viable CD45^+^NK1.1^+^ cells (**Fig. 1B**, **Fig. S1A**). In total, 11,978 NK1.1^+^ cells passed quality-control filtering and were retained for analysis (**Fig. S1B-C**).

Integrated analysis of brain- and liver-derived cells identified three major clusters: NK cells, ILC1s, and T cells (**Fig. S1B**). The T cell cluster, characterized by high *Cd3e* and low *Ncr1* (NKp46) expression, was excluded from further analysis (**Fig. S1D, Supplementary Table 1**). Re-clustering of the remaining 10,291 cells resolved two principal populations: NK cells, marked by *Itga2* and *Eomes*, and ILC1s, marked by *Itga1*, *Cd200r1*, and *Zfp683* (**Fig. 1C-D**).

Subset analysis identified six populations of NK cells and three ILC1 cell states based on canonical markers and top differentially expressed genes (**Fig. 1C**, **Fig. S2A-B, Supplementary Table 2**). As expected, ILC1s scored highly for genes associated with tissue residency (*Cd69*, *Cxcr6*, *Zfp683*), whereas NK cells scored highly for circulating marker genes (*S1pr5*, *Klf2*, *Sell*) (**Fig. S3A, Supplementary Tables 3-5**)^43^. The six NK cell states differed in their expression of markers associated with maturity and activation. We identified two mature NK cell states, which we termed “circulating NKs” and “NF-κB-activated NKs.” Both express Ly49-family receptors (including *Klra8*, *Klra4*, and *Klra9*), cytokine-response and trafficking receptors (*Il18r1*, *Il12rb2*, and *Cx3cr1*), and adhesion-associated molecules such as *Itgam* and *Spn* (**Fig. 1E**, **Fig. S3B**). Both states also express high levels of *Gzma* and *Prf1*, consistent with cytotoxic effector potential. “Circulating NKs” show the highest circulation score of all NK subsets (**Fig. S3A**), suggesting these cells may have been extracted from capillaries within the tissues. The “NF-κB-activated NK” state was further distinguished by increased expression of NF-κB pathway-associated genes^44,45^, which is indicative of heightened activation, together with *Ccl3* and *Ccl4*, chemokines commonly induced in activated NK cells that can promote recruitment of additional immune cells (**Fig. 1E**, **Fig. S3C, Supplementary Tables 3-5**)^46,47^. To rule out artifactual activation during tissue processing, we calculated a stress/immediate-early gene score, which did not differ between NF-κB-activated NKs and the other NK states, supporting the conclusion that this program was not an artifact (**Fig. S3D, Supplementary Tables 3-5**). An “immature NK” state was defined by *Cd27*, *Ctla2a*, and enrichment of ribosomal transcripts, consistent with a less differentiated, biosynthetically active phenotype (**Fig. 1E**, **Fig. S2A**)^48,49^. A smaller cluster characterized by *Ifng* together with the interferon-stimulated genes (ISGs) *Isg15*, *Ifit3*, *Bst2*, *Stat1*, and *Zbp1* was annotated as “ISG+ NKs” (**Fig. 1E**). Previous work has shown that ISG-high NK cells represent a transient activation state that results from acute interferon exposure^50^. We also identified a population of “pre-NKs” that express *Nme1*, *Nme2*, *Eif5a*, and *Ran*, suggesting a precursor-like, proliferative program. Finally, we resolved a population of “cycling NKs” that express canonical cell-cycle genes (*Stmn1*, *Hmgb2*, *Pclaf*, *Tubb5*, *Dut*, and *Cks1b)*.

Within the ILC1 compartment, the three cell states also appear to segregate based on maturation state. We found a “progenitor-like ILC1” population expressing *Rora*, *Il7r*, and *Emb*, but lacking expression of cytotoxic effector genes such as *Gzmb* and *Gzmc* (**Fig. 1E**, **Fig. S3E-F**)^51^. “Cytotoxic ILC1s” and “immature-like ILC1s” share a partially overlapping signature that includes *Cd7*, *Rgs1*, *S100a4*, and *Xcl1*. However, “Cytotoxic ILC1s” express higher levels of *Gzmb*, consistent with greater cytotoxic potential, whereas “immature-like ILC1s” are distinguished by increased expression of *Smyd3*, *Mef2a*, *Lgals3*, *Rps15a*, and *Rpl22*, as well as genes related to the immature NK cell cluster.

Remarkably, we find that the composition of NK and ILC1 populations is strikingly different between the brain and liver. The liver exhibits greater overall group 1 ILC diversity, with large frequencies of the most activated effector NK (30% NF-κB-activated) and ILC1 (30% cytotoxic ILC1) subsets (**Fig. 1F-G**). In contrast, the brain contains rare populations of both of these cell states (3.3% and 0.6%, respectively). The predominant population captured from the brain was “circulating NK” cells, which represent 65% of all group 1 ILCs in brain tissue (**Fig. 1F-G**), suggesting that the majority of cells captured from the brain may have localized within capillaries. We recovered small frequencies of “immature NK (15%)” and “pre-NK” (5%), suggesting that small numbers of less activated NK cells may be found in the brain parenchyma. Interestingly, we also found a small population of “immature-like ILC1s” (7%), showing that the brain does contain resident ILC1s, but their activation state is diminished relative to the “cytotoxic ILC1s” found in the liver. Importantly, the subset distributions were consistent across individual mice (**Fig. S3G**). Notably, we found limited differences between metastatic and control conditions in the liver and brain, suggesting that the response to tumor growth does not shift subset frequencies. Together, these data show that the brain contains substantially restricted group 1 ILC diversity compared to the liver, with notably reduced effector NK and ILC1 cell populations and a preponderance of circulating NK cells, raising the question of whether NK cells penetrate metastatic lesions in the CNS.

### NK cells fail to infiltrate BCBM

To investigate group 1 ILC-cell localization within the brain parenchyma, we assessed the intravascular versus extravascular distribution of NK cells and ILC1s using intravascular antibody staining. Mice were injected i.v. with PE-Cy5-conjugated anti-CD45 antibody three minutes before euthanasia (**Fig. 2A**). Following organ harvest and tissue digestion, isolated cells were stained *ex vivo* with a second, Pacific Blue-conjugated anti-CD45 antibody. This approach allows discrimination of intravascular leukocytes, which are labeled by both antibodies, from extravascular leukocytes that are inaccessible to the i.v. administered antibody and therefore stained only with the *ex vivo* anti-CD45 antibody. Intravascular anti-CD45 staining has been validated as a method to distinguish vascular from tissue-localized leukocytes, with rapid labeling of the vast majority of blood-accessible leukocytes within minutes of antibody administration^52^. We applied this approach to healthy control mice and to mice at 4 or 10 days after intracardiac injection of EO771 cells. As expected, this assay was not informative in the liver because of its highly permeable vasculature, with nearly all NK1.1^+^ cells staining positive for the i.v.-administered CD45 antibody even within this short labeling period (**Fig. S4A-B**). In contrast, in the brain, which is protected by the tightly regulated BBB, we readily distinguish i.v. CD45^hi^ intravascular and i.v. CD45^lo^ extravascular NK1.1^+^ populations (**Fig. 2B**, **Fig. S4B**). Analysis of CD49a and CD49b expression showed that nearly all intravascular NK1.1^+^ cells in the brains of control and tumor-bearing animals were CD49b^+^ NK cells (88%), whereas the extravascular compartment was composed predominantly of CD49a^+^ ILC1s (63%) and a small population of CD49a^+^CD49b^+^ ILC1-like cells (18%) (**Fig. 2C**, **Fig. S4C**). Interestingly, in mice with brain metastasis, we also observed a small population of extravascular CD49b^+^ NK cells not present in control animals. These findings support the conclusion from our scRNA-seq data that most CD49b^+^ NK cells detected in the brain correspond to a circulating population extracted from blood vessels, and that small numbers of these cells enter the brain parenchyma in response to tumor growth (**Fig. 1F-G**). Quantification of total cell numbers revealed substantially lower NK1.1^+^ cell infiltration in the brain than in the liver (**Fig. 2D**). In the brain, metastasis was associated with only a modest increase in the total number of extravascular ILC1s and NKs, further supporting that there is a response to tumor growth, but that tissue infiltration is limited (**Fig. 2E**). By contrast, the number of intravascular NKs in the brain remained unchanged during metastatic progression (**Fig. S4D**). In the liver, total numbers of ILC1s, ILC1-like cells, and NKs were not altered after EO771 injection, suggesting that liver-resident group 1 ILC populations control metastatic seeding without requiring major expansion (**Fig. S4E**). Together, these data show that brain metastasis invokes a response from both tissue-resident ILC1s and infiltrating NK cells, but it is substantially more limited than in the liver.

**Figure 2:**
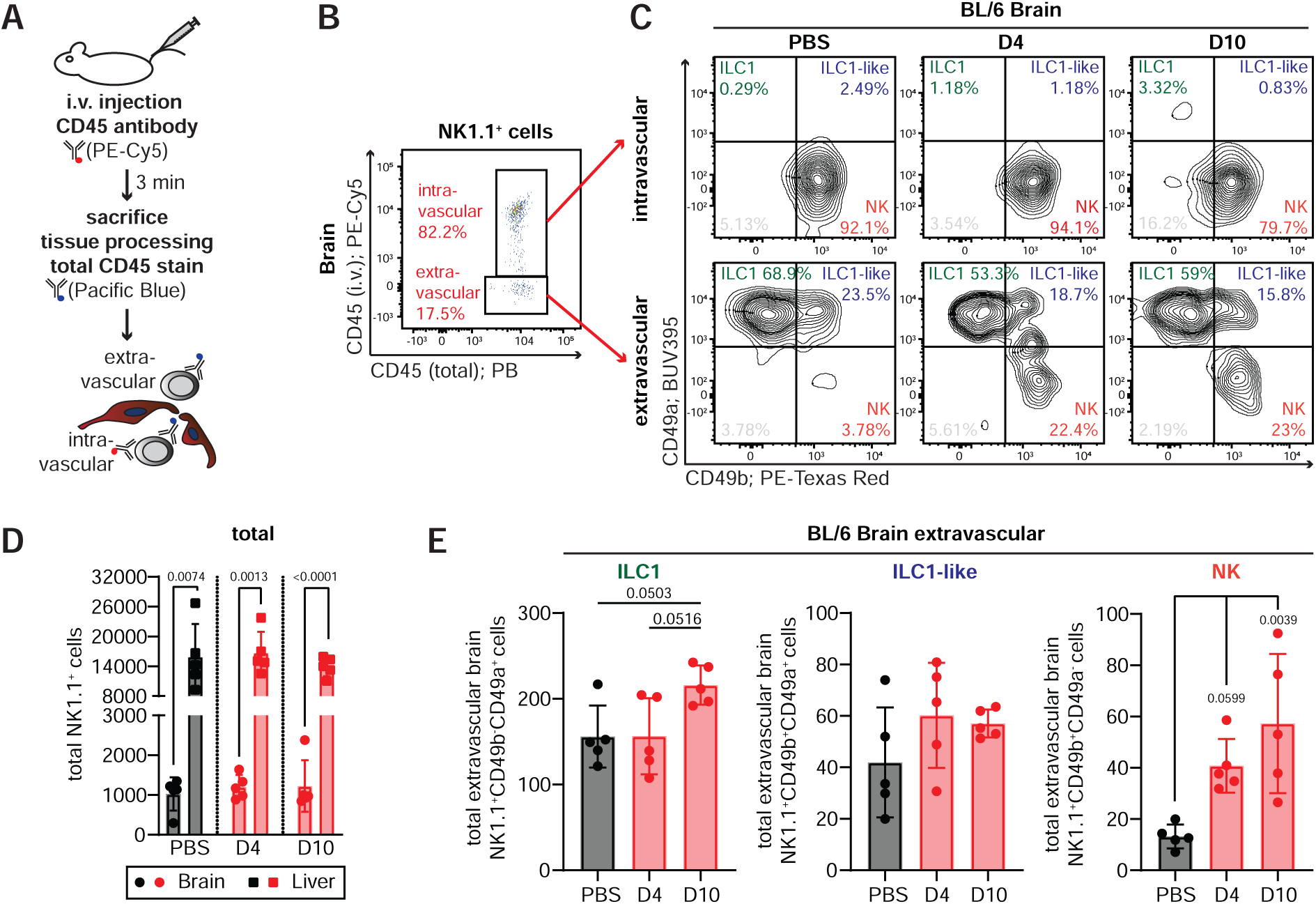
NK cells fail to infiltrate BCBM. **(A)** Schematic of how the intravascular CD45 staining works. Mice are injected (i.v.) with a PE-Cy5-conjugated CD45 antibody 3 minutes before sacrifice. After tissue processing, we stain total CD45 with a Pacific Blue-conjugated antibody. Using flow cytometry, we can then distinguish between PE-Cy5/Pacific Blue double-positive intravascular immune cells and Pacific Blue single-positive extravascular immune cells. **(B)** Flow cytometry gating strategy to identify intravascular (CD45 i.v.^+^/CD45 total^+^) and extravascular (CD45 i.v.^−^/CD45 total^+^) NK1.1^+^ cells in murine brain. Cells were pregated on viable, lineage^-^ (lin^-^; CD19^-^Gr-1^-^Ter119^-^EpCAM^-^CD4^-^CD8^-^), CD45^+^ cells. **(C)** Flow cytometry gating strategy to define brain-resident CD49a^+^ ILC1s, CD49a^+^CD49b^+^ ILC1-like cells, and CD49b^+^ NKs in PBS- or EO771 eGfly-injected mice. Top: intravascular NK1.1^+^ gate; bottom: extravascular NK1.1^+^ gate. **(D)** Absolute counts of total NK1.1^+^ cells in brains and livers from PBS (black)- or EO771 eGfly-injected (red) mice (n=5 per group and organ). **(E)** Absolute counts of extravascular ILC1s, ILC1-like cells, and NKs in brains of PBS-(black) or EO771 eGfly-injected mice (red; n=5 per group and organ). Data are represented as mean ± SD. Statistics: two-tailed Student’s t-test (D) and one-way ANOVA (E). Exact p-values are shown for statistically significant comparisons.

### Group 1 ILCs fail to control brain metastasis

Next, we investigated the functional capacity of the group 1 ILC response to control brain metastasis in the E0771 intracardiac model *in vivo*. We detected a higher metastatic burden in the brain than in the liver of BL/6 mice 4 and 10 days after tumor cell inoculation by *ex vivo* bioluminescence imaging (BLI) (**Fig. S5A-B**). This enhanced control of liver metastasis is independent of adaptive immunity, as Rag1^-/-^ mice also display a lower liver metastatic burden than in the brain (**Fig. S5C-D**). Given that NK cells and ILC1s are required to control liver metastasis and are the only cytotoxic lymphocyte populations present in Rag1^-/-^ mice, we hypothesized that differential control of metastasis in the liver and brain reflects tissue-specific phenotypic and functional differences in group 1 ILCs.

To test this hypothesis, we performed systemic depletion of group 1 ILCs in immunocompetent BL/6 mice using an antibody against NK1.1, followed by intracardiac injection of EO771 cells (**Fig. 3A**). The anti-NK1.1 antibody efficiently depletes group 1 ILCs in the liver and brain (**Fig. 3B**). While the tumor burden in the body of mice depleted of NK1.1^+^ cells was significantly more severe compared to control animals, the mice in both groups displayed indistinguishable *in vivo* tumor outgrowth in the head region (**Fig. S5E**). *Ex vivo* analysis using BLI and flow cytometry demonstrated that systemic depletion of NK1.1^+^ cells did not affect brain metastasis, even as liver metastasis became substantially more severe (**Fig. 3C-F**). These results imply that NK1.1^+^ cells have a limited intrinsic capacity to control BCBM.

**Figure 3.**
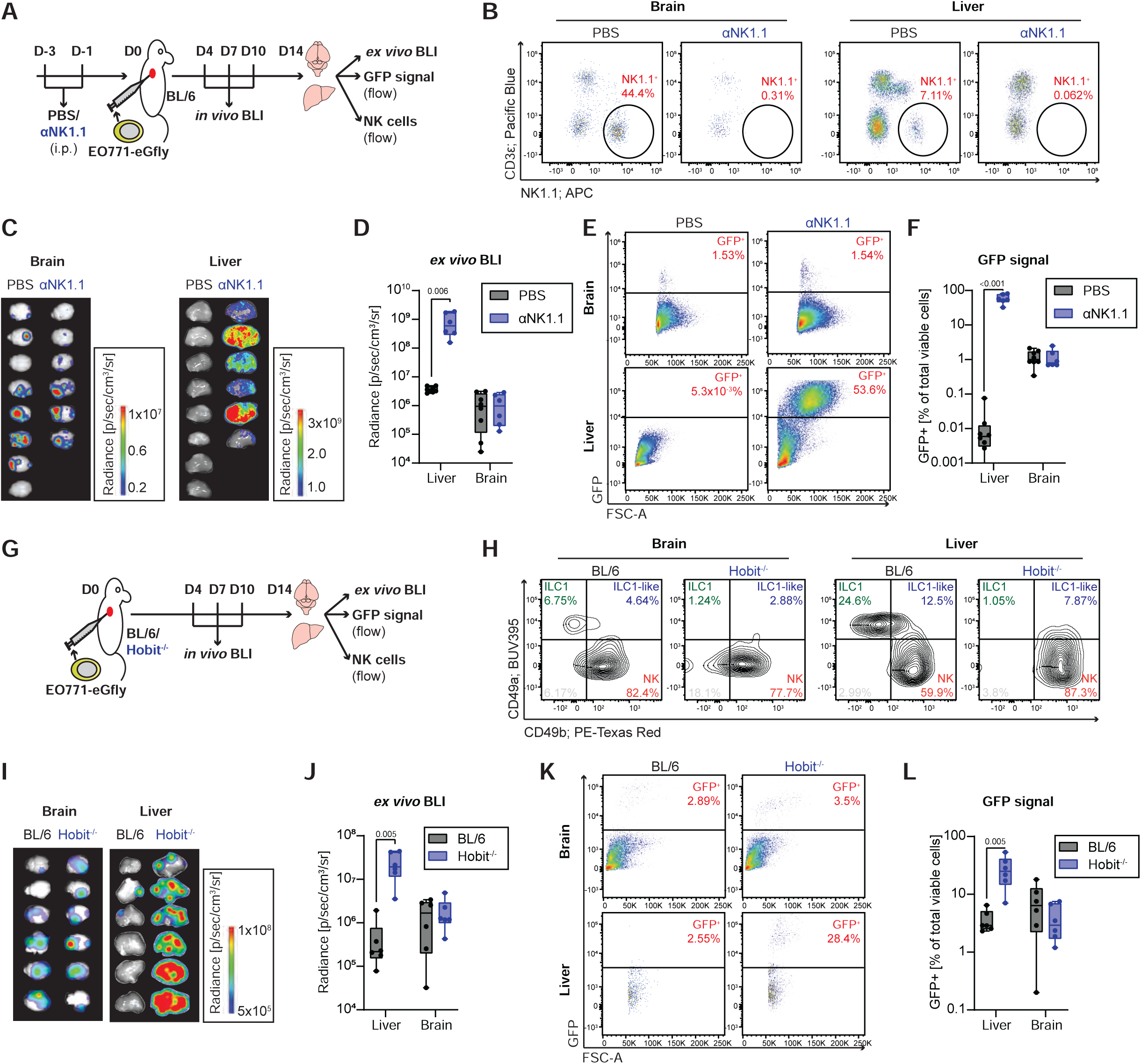
Group 1 ILCs fail to control brain metastasis. **(A)** Experimental schematic for systemic NK1.1^+^ cell depletion. BL/6 mice received an anti-NK1.1 depleting antibody (αNK1.1) 3 and 1 days prior to intracardiac injection of EO771 eGFP/luciferase (eGfly) cells; control mice received PBS instead of αNK1.1. Tumor burden was monitored by bioluminescence imaging (BLI) and flow cytometry, and NK1.1^+^ cell abundance was assessed at the experimental endpoint. **(B)** Flow cytometry gating strategy for NK1.1^+^ cells (NK1.1^+^CD3ε^−^) in brains and livers of control and NK1.1-depleted mice. Cells were pre-gated on viable CD45^+^ leukocytes. **(C-D)** *Ex vivo* BLI of brains and livers harvested on day 14 post-injection (C) and corresponding quantification of total bioluminescence signal (D) from control (PBS; n=8) and NK1.1-depleted (αNK1.1; n=6) mice. **(E-F)** Flow cytometry gating strategy (E) and quantification (F) of GFP^+^ EO771 eGfly tumor cells in digested brains and livers from control (PBS; n=8) and NK1.1-depleted (αNK1.1; n=6) BL/6 mice. Cells were pre-gated on viable cells. **(G)** Experimental schematic for the Hobit-knockout (Hobit^-/-^) experiment. BL/6 or Hobit^-/-^ mice were intracardially injected with EO771 eGfly cells. Tumor burden was monitored by bioluminescence imaging (BLI) and flow cytometry, and the abundance of various NK1.1^+^ subsets was assessed at the experimental endpoint. **(H)** Flow cytometry gating strategy to define CD49a^+^ ILC1s, CD49a^+^CD49b^+^ ILC1-like cells, and CD49b^+^ NKs in the brains and livers of EO771 eGfly-injected BL/6 and Hobit^-/-^ mice. Cells were pregated on viable, lineage-negative (lin^-^; CD19^-^Gr-1^-^Ter119^-^EpCAM^-^CD4^-^CD8^-^), CD45^+^NK1.1^+^ cells. **(I-J)** *Ex vivo* BLI of brains and livers harvested on day 14 post-injection (I) and corresponding quantification of total bioluminescence signal (J) from BL/6 (n=7) and Hobit^-/-^ (n=7) mice. **(K-L)** Flow cytometry gating strategy (K) and quantification (L) of GFP^+^ EO771 eGfly tumor cells in digested brains and livers from BL/6 (n=7) and Hobit^-/-^ (n=7) mice. Cells were pre-gated on viable cells. Data are represented as mean ± SD. Statistics: two-tailed Student’s t-test (D, F, J, L). Exact p-values are shown for statistically significant comparisons.

To test the role of the tissue-resident ILC1s in brain metastasis, we performed intracardiac injections of EO771 cells in mice deficient for Hobit (Hobit^-/-^ mice), a transcription factor required for ILC1 development and tissue residency (**Fig. 3G**). Hobit^-/-^ mice had drastically reduced ILC1 counts in the liver and brain compared to BL/6 controls (**Fig. 3H)**. Longitudinal *in vivo* imaging revealed that Hobit^-/-^ mice had more severe metastasis in the body region than BL/6 controls by day 10, while no differences in metastatic burden in the head region were detectable (**Fig. S5F**). *Ex vivo* BLI and flow cytometry analysis did not show a significant difference in tumor burden in the brains of Hobit^-/-^ mice compared with controls, whereas Hobit deficiency induced more severe liver metastasis (**Fig. 3I-L**). On this basis, we conclude that ILC1s do not contribute to the control of BCBM under normal physiological conditions.

In addition to group 1 ILCs, components of the adaptive immune system contribute to cytotoxicity-mediated control of metastasis. We hypothesized that depleting group 1 ILCs in our previous experiments may have led to compensatory anti-metastatic immunity driven by the adaptive immune system, thereby masking the functional contributions of group 1 ILCs to metastasis control. To test this hypothesis, we depleted group 1 ILCs in T/B cell-deficient Rag1^-/-^ mice using the anti-NK1.1 antibody (**Fig. S5G**). We then challenged NK1.1-depleted mice and matched controls with intracardiac injection of EO771 tumor cells and assessed brain metastasis outgrowth and immune response. Confirming our findings in BL/6 mice, NK1.1 depletion increased metastasis burden in peripheral tissues (**Fig. S5H**). In contrast to our findings in BL/6 mice, Rag1^-/-^ mice had more severe metastatic disease in the head region after NK1.1 depletion. *Ex vivo* BLI and flow cytometry assessment of tumor burden confirmed that group 1 ILC depletion in Rag1^-/-^ mice exacerbates both liver and brain metastasis (**Fig. S5I-L**). In summary, we demonstrate that group 1 ILCs can control both liver and, partially, brain metastasis in the absence of adaptive immunity. This is consistent with our scRNA-seq and flow cytometry data, which showed a modest increase in group 1 ILCs in the brain during metastasis formation. However, these results led us to hypothesize that group 1 ILCs differ functionally and phenotypically between the liver and brain, explaining their lack of full anti-metastatic function in the brain.

### Limited IL15/IL-15Rα expression in the CNS

NK cells and ILC1s depend on continuous cytokine signals for their maintenance, survival, and functional competence within tissues^31^. Given our finding that group 1 ILCs fail to efficiently infiltrate or control metastatic growth in the brain, whereas they restrict metastasis formation in the liver, we hypothesized that these tissue-specific differences may partly reflect differences in the local availability of signals that recruit, maintain, or activate group 1 ILCs. We therefore analyzed publicly available mouse brain and liver datasets for the expression of chemokines and cytokines implicated in group 1 ILC recruitment and regulation. The Mouse Brain Immune Atlas revealed expression of several NK-cell-recruiting chemokines, including *Ccl3*, *Ccl4*, and *Ccl5*, predominantly among border-associated macrophages, dendritic cells, NK cells, and microglia (**Fig. 4A**; **Fig. S6A**). These populations also expressed inflammatory cytokines such as *Il1b*, whereas the inhibitory cytokine *Tgfb1* was broadly expressed across brain immune populations. In the liver, several immune populations expressed *Ccl3* and *Ccl4*, including basophils, ILC1s, NK cells, and dendritic cells (**Fig. 4B**; **Fig. S6B**). *Il1b* expression was prominent in neutrophils, monocytes/macrophages, dendritic cells, and Kupffer cells, while *Tgfb1* was broadly expressed.

**Figure 4.**
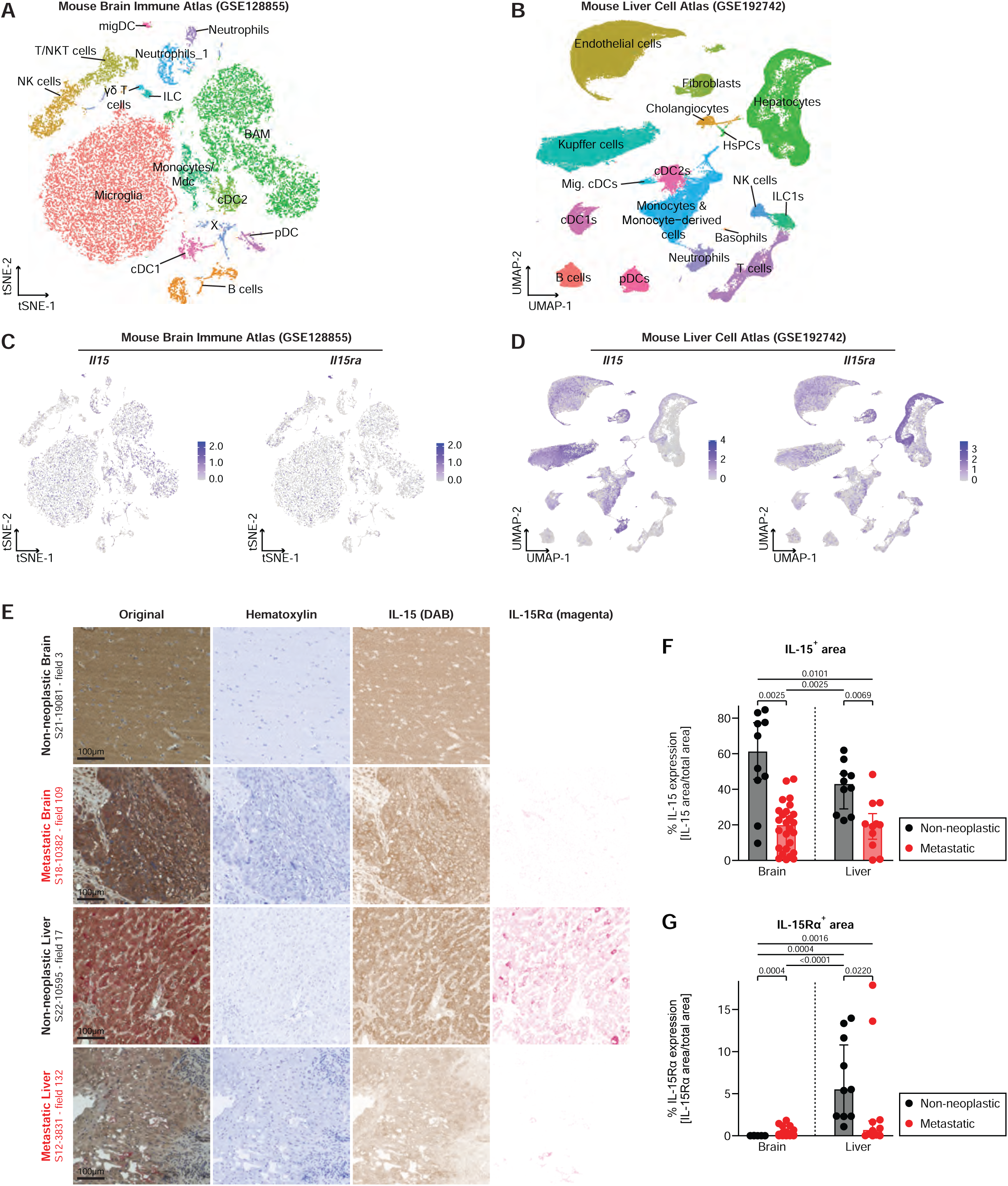
Limited IL15/IL-15Rα expression in the CNS. **(A)** tSNE visualization of scRNA-seq profiles from all cells of the Mouse Brain Immune Atlas (GSE12885), colored by main cell types. migDC: migratory dendritic cells; BAM: border-associated macrophages; cDC: conventional dendritic cells; pDCs: plasmacytoid dendritic cells. **(B)** UMAP visualization of scRNA-seq profiles from all cells of the Mouse Liver Cell Atlas (GSE192742), colored by main cell types. HsPCs: hematopoietic stem and progenitor cells; Mig. cDCs: migratory conventional dendritic cells; cDCs: conventional dendritic cells; pDCs: plasmacytoid dendritic cells. **(C-D)** Feature plots showing relative expression of *Il15* and *Il15ra* in the Mouse Brain Immune Atlas (C) and the Mouse Liver Cell Atlas (D). **(E-G)** *In situ* IL-15 and IL-15Rɑ protein expression analyzed by immunohistochemistry in a cohort of 57 human brain and liver samples (10 non-neoplastic brain sections, 26 metastatic brain sections, 10 non-neoplastic liver sections, 11 metastatic liver sections). Micrographs show representative FOVs with complete staining (original) or after deconvolution into the hematoxylin, IL-15 (DAB), and IL-15Rɑ (magenta) channels (E). Scale bar: 100 µm. Quantification of the IL-15-positive (F) and IL-15-Rɑ-positive (G) areas in each section, shown as the frequency of the total area. Data are represented as mean ± SD. Statistics: Kruskal–Wallis across the four groups, followed by two-sided Mann–Whitney *U* tests for all six pairwise comparisons with Benjamini–Hochberg false-discovery-rate correction (F, G). Exact p-values are shown for statistically significant comparisons.

We next focused on the IL-15–IL-15Rα axis, which is critical for group 1 ILC homeostasis and activation. Both *Il15* and *Il15ra* expression were very limited among brain immune populations, whereas several hepatic populations expressed components of this pathway (**Fig. 4C–D**). Monocytes/macrophages expressed *Il15*, while hepatocytes, endothelial cells, and Kupffer cells showed prominent *Il15ra* expression with detectable *Il15*. Consistent with these transcriptomic data, IL-15Rα protein was readily detectable in murine liver but was little to undetectable in brain lysates by immunoblotting (**Fig. S6C**). IL-15 is predominantly delivered to NK cells and ILC1s through trans-presentation, whereby IL-15 bound to IL-15Rα on neighboring cells engages IL-2/15Rβ and the common γ-chain on responding group 1 ILCs^53,54^. Thus, the abundance of IL-15Rα-expressing cells can critically determine local IL-15 bioavailability, even in the presence of IL-15.

To determine whether this tissue-specific difference was conserved in humans, we analyzed IL-15 and IL-15Rα protein expression in non-neoplastic and metastatic human brain and liver tissues using duplex immunohistochemistry (n=10 non-neoplastic brain samples, n=26 metastatic brain samples, n=10 non-neoplastic liver samples, n=11 metastatic liver samples; **Supplementary Table 6**). We quantified staining using a standardized image-analysis pipeline incorporating tissue segmentation, predefined field-of-view sampling, chromogen deconvolution, quality-control exclusion criteria, and experimentally validated staining thresholds (**Fig. S7**). IL-15 staining was abundant in both non-neoplastic brain and liver and decreased in metastatic tissue in both organs (**Fig. 4E–F**; **Fig. S8–S9**). In striking contrast, IL-15Rα-positive area was approximately 500-fold lower in non-neoplastic brain than in non-neoplastic liver (**Fig. 4G**). IL-15Rα staining increased in metastatic brain tissue but remained markedly below hepatic levels, whereas liver metastases showed a reduction relative to non-neoplastic liver.

Together, these data identify a pronounced tissue-specific difference in the IL-15/IL-15Rα axis: although IL-15 protein is abundant in both brain and liver, IL-15Rα is markedly restricted in the brain. Because IL-15Rα-mediated trans-presentation is a major mechanism supporting NK-cell survival and function, limited IL-15 trans-presentation may contribute to the impaired group 1 ILC response to brain metastases.

### IL-15/IL15Ra rescues group 1 ILC diversity and expands effector populations in BCBM

Because NK cells and ILC1s depend on IL-15/IL-15Rɑ for proliferation and tissue persistence, particularly within the tumor microenvironment, and because IL-15 has been shown to enhance NK cell anti-tumor activity, we hypothesized that therapeutic IL-15/IL-15Rɑ supplementation could improve control of brain metastasis by augmenting group 1 ILC responses in the brains of immunocompetent mice^55–58^. To test this, we treated mice bearing brain metastases with an IL-15 superagonist consisting of soluble IL-15 complexed with soluble IL-15 receptor α (IL-15Rα), hereafter referred to as IL-15 complex (IL-15c)^59–61^.

To determine whether exogenous IL-15c alters NK cell populations at the transcriptional level in the brains of metastasis-bearing mice, we performed transcranial co-injections of IL-15c and EO771 tumor cells, followed by an additional dose of IL-15c on day 1 after the initial injection (**Fig. 5A**). Mice were euthanized on day 4, and brains were harvested, confirming comparable tumor cell engraftment between groups (**Fig. S10A**). Brain leukocytes were isolated, enriched for NK1.1^+^ group 1 ILCs using MACS and FACS, and subjected to single-cell RNA sequencing. After quality control, we recovered 7,057 NK1.1+ cells, of which 1,126 NKT cells were excluded (**Fig. S10B–D, Supplementary Table 7**). The remaining 5,931 group 1 ILCs were integrated with 10,291 cells from the time-course experiment (**Fig. 1C**), generating an “All Cells” object containing 16,222 cells (**Fig. 5B**). Clustering quality remained high after integration, and shared marker genes were consistently identified across both datasets (**Fig. S11A-B, Supplementary Table 8**). IL-15c treatment profoundly reshaped the composition of NK cell and ILC1 subclusters in the brain, generating a phenotypic diversity that closely resembled the distribution of group 1 ILCs in the liver (**Fig. 5C**). Specifically, the predominance of circulating NK cells in the brain was reduced, accompanied by a relative expansion of immature NK cells and NF-κB NK cells (**Fig. 5D**). Immature-like ILC1s also expanded markedly following IL-15c treatment. In addition, the increased frequency of cycling NK cells suggests that IL-15c promotes local proliferation of group 1 ILCs within the brain. Together, these findings indicate that IL-15c drives both diversification and proliferation of group 1 ILCs in metastatic brains.

**Figure 5.**
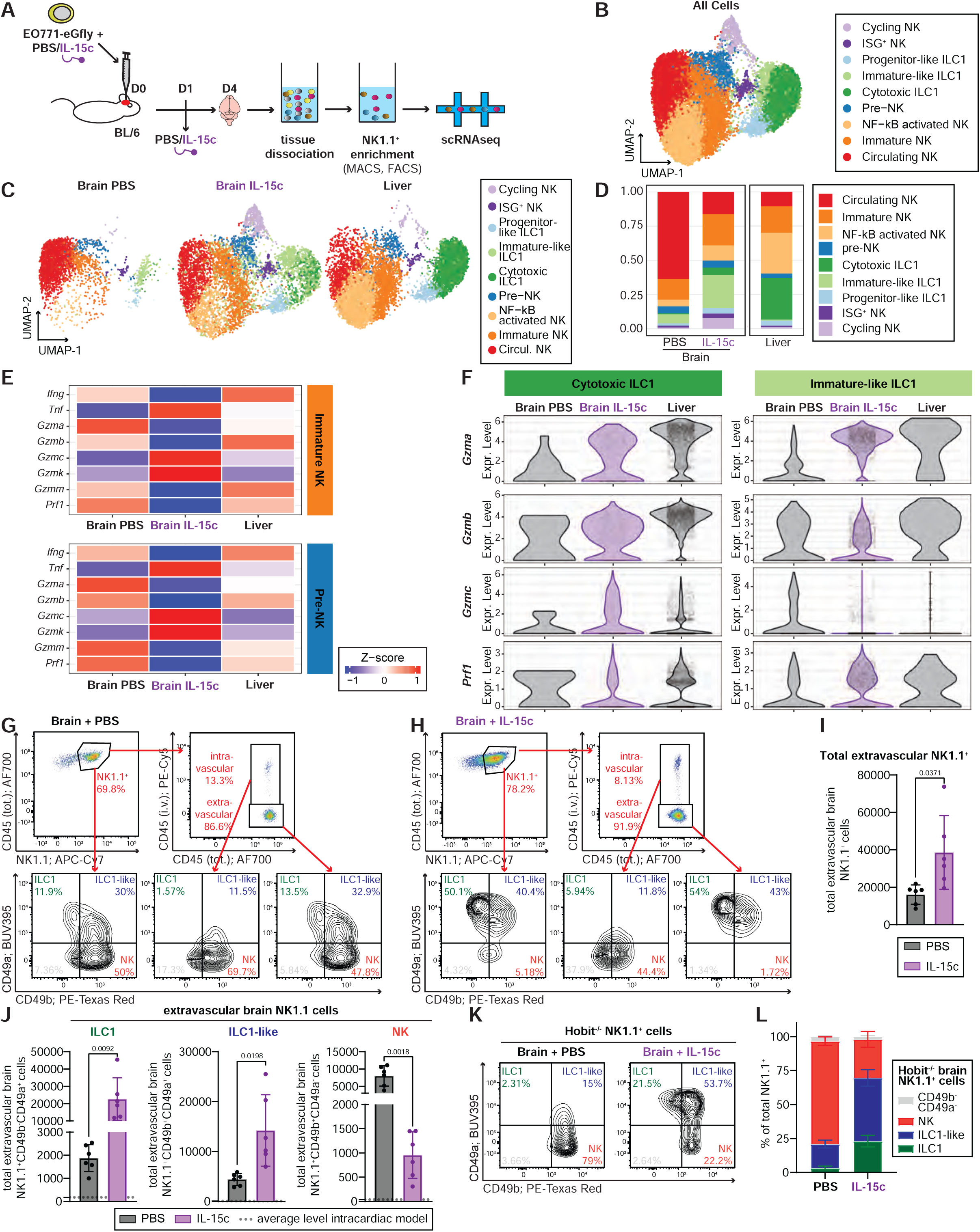
IL-15/IL15Ra rescues group 1 ILC diversity and expands effector populations in BCBM. **(A)** Experimental workflow for isolation and sequencing of metastasis-infiltrating NK1.1^+^ group 1 ILCs from brains of PBS- and IL-15 complex (IL-15c)-treated mice. BL/6 mice received intracranial EO771 eGfly cells together with PBS (n=5) or IL-15c (n=5). Mice received an additional dose of PBS or IL-15c on day 1 post-implantation and were euthanized on day 4. Brains were enzymatically digested, and NK1.1^+^ cells were enriched by magnetic-activated cell separation (MACS) followed by fluorescence-activated cell sorting (FACS) prior to droplet-based scRNA-seq. **(B–C)** UMAP visualization of scRNA-seq profiles from sorted NK1.1^+^ cells isolated from PBS- and IL-15c-treated brains, together with NK1.1^+^ cells from livers of PBS-injected (healthy) and tumor-burdened BL/6 mice (from Fig. 1). (B) shows all cells in a single embedding, and (C) shows cells split by tissue of origin and treatment condition. **(D)** Stacked bar plot showing the proportional representation of each transcriptional cluster among NK1.1^+^ cells from PBS-treated brains, IL-15c-treated brains, and liver samples. **(E)** Heat map of Z-scored expression of NK-associated effector granzymes and cytokines in the “Immature NK” and “Pre-NK” cell clusters, split into the Brain PBS, Brain IL-15c, and Liver groups. **(F)** Violin plots showing the expression of *Gzma*, *Gzmb*, *Gzmc*, and *Prf1* in cytotoxic ILC1s or immature-like ILC1s, split by tissue and treatment. **(G-H)** Flow cytometry gating strategy to identify intra- and extravascular brain CD49a^+^ ILC1s, CD49a^+^CD49b^+^ ILC1-like cells, and CD49b^+^ NKs in EO771 eGfly–injected mice treated with PBS (G) or IL-15c (H). Mice received three doses of PBS or IL-15c (D0, D1, and D4) and were sacrificed on day 7. **(I-J)** Absolute numbers of total extravascular NK1.1^+^ (I), ILC1s, ILC1-like cells, and NKs (J) in brains of BL/6 mice that were intracranially injected with EO771 eGfly cells, in combination with PBS (n=6) or IL-15c (n=7). Mice received three doses of PBS or IL-15c (D0, D1, and D4) and were sacrificed on day 7. Dotted lines indicate the mean cell counts for the corresponding NK1.1^+^ subsets in BL/6 mice following intracardiac EO771 eGfly injection. **(K)** Flow cytometry gating strategy to identify brain CD49a^+^ ILC1s, CD49a^+^CD49b^+^ ILC1-like cells, and CD49b^+^ NKs in EO771 eGfly–injected Hobit^-/-^ mice treated with PBS or IL-15c. Mice received three doses of PBS or IL-15c (D0, D1, and D4) and were sacrificed on day 7. **(L)** Bar plot representing the frequency of brain CD49a^+^ ILC1s, CD49a^+^CD49b^+^ ILC1-like cells, and CD49b^+^ NKs in EO771 eGfly–injected Hobit^-/-^ mice treated with PBS (n=6) or IL-15c (n=6). Data are represented as mean ± SD. Statistics: two-tailed Student’s t-test (I, J). Exact p-values are shown for statistically significant comparisons.

We next examined the expression of canonical NK cell and ILC1-associated genes. Notably, immature NK and pre-NK cells from brains of IL-15c-treated mice showed increased expression of *Itga1* (CD49a), suggesting acquisition of a more tissue-adapted phenotype (**Fig. S12A**)^22,23^. IL-15c treatment was also associated with induction of an effector program in these two clusters characterized by increased expression of *Tnf*, *Gzmc*, and *Gzmk* (**Fig. 5E**). This is consistent with previous studies showing that IL-15 stimulation induces CD49a and granzyme expression in NK cells, including rapid induction of Granzyme B followed by delayed induction of Granzyme C^62,63^. In parallel, IL-15c increased the expression of several activating and inhibitory receptor-associated genes, including *Cd28*, *Tgfbr1*, *Tgfbr2*, *Klrg1*, and *Havcr2* (**Fig. S12B-C**). Together, these findings indicate that intracranial IL-15c treatment promotes a tissue-adapted and effector-associated NK-cell state characterized by CD49a induction, expression of a Granzyme C-centered effector program, and simultaneous upregulation of regulatory receptor pathways. These data are consistent with the emergence of a highly activated, but tightly regulated, tissue-infiltrating NK-cell population in the brain.

Although the immature-like ILC1s in the brain of IL-15c-treated mice remained transcriptionally distinct from the cytotoxic ILC1 state, IL-15c induced a more effector-like phenotype in these cells, marked by increased expression of *Gzma* and *Prf1* (**Fig. 5F**). Collectively, these data show that IL-15c treatment enhances group 1 ILC diversity and proliferation in metastatic brains, promotes the transition of NK cells toward tissue residency, and augments the cytotoxic potential of tissue-resident ILC1s. These effects likely contribute to the anti-metastatic activity of IL-15c.

Having established that IL-15c enriches disease-responsive group 1 ILC subpopulations, we next sought to determine whether these cells extravasate from the circulation into the brain parenchyma to mediate tumor control. To address this, we performed flow cytometric analysis of metastasis-infiltrating group 1 ILCs in IL-15c-treated brains using an intravenously injected CD45 antibody to distinguish intravascular from extravascular cells (**Fig. 2A**). BL/6 mice were intracranially co-injected with IL-15c and EO771 tumor cells on day 0, followed by intracranial booster doses of IL-15c on days 1 and 4. On day 7, mice were euthanized, and brain group 1 ILCs were isolated and analyzed by flow cytometry. In both PBS- and IL-15c-treated mice, most NK1.1^+^ cells were not labeled by the intravenously administered CD45 antibody and were therefore classified as extravascular (**Fig. 5G-H**). This contrasts sharply with the intracardiac model, in which only 20% of NK1.1^+^ cells were extravascular (**Fig. 2B**). This difference is likely explained by the inoculation route: intracranial injection may disrupt the brain vasculature, thereby facilitating immune cell access to the parenchyma, and may also generate larger brain metastases that promote immune infiltration. In PBS-treated brains, most NK1.1^+^ cells in both the intravascular and extravascular compartments displayed a CD49b^+^ NK cell phenotype (**Fig. 5G**). In contrast, IL-15c treatment shifted the NK1.1^+^ compartment toward CD49a^+^ ILC1s and CD49a^+^CD49b^+^ ILC1-like cells (**Fig. 5H**). IL-15c also increased the total number of extravascular NK1.1^+^ cells, driven by higher numbers of ILC1s and ILC1-like cells and accompanied by a reduction in NK cell numbers (**Fig. 5I–J**). Overall, the number of extravascular NK1.1^+^ cells was substantially higher in this model than in the intracardiac setting. We repeated this experiment in Rag1^-/-^ mice and observed the same increase in extravascular ILC1s and ILC1-like cells, both in frequency and absolute number (**Fig. S13A–D**). Finally, we examined the group 1 ILC landscape in the brains of Hobit^-/-^ mice following IL-15c treatment. Similar to BL/6 and Rag1^-/-^ mice, IL-15c induced the accumulation of ILC1-like cells in Hobit^-/-^ mice, along with a partial increase in ILC1s (**Fig. 5K-L**). These findings suggest that NK1.1^+^ cells retain some plasticity, allowing infiltrating NK cells to partially compensate for the absence of tissue-resident ILC1s in Hobit^- /-^ mice. Furthermore, it indicates that the CD49b^+^CD49a^+^ NK cells originate from CD49b^+^ NK cells. Overall, these findings demonstrate that IL-15c reprograms the metastatic brain group 1 ILC compartment by promoting extravascular accumulation, phenotypic diversification, tissue residency, and cytotoxic function.

### Group 1 ILCs are critical to limit brain metastasis after IL-15c treatment

BL/6 mice received intracranial co-injections of EO771 tumor cells together with IL-15c, followed by two additional intracranial IL-15c injections on days 1 and 4 after tumor inoculation (**Fig. 6A**). By day 7, IL-15c-treated BL/6 mice showed significantly improved tumor control compared with PBS controls (**Fig. 6B-C**). This anti-tumor effect is independent of ILC1s, as Hobit^-/-^ mice showed a similar response (**Fig. 6D-E**). Repeating the experiment in Rag1^-/-^ mice likewise showed improved tumor control after IL-15c treatment, indicating that the protective effect is independent of T and B cells (**Fig. 6F-G**). To determine whether NK cells mediate the protective effect of IL-15c, we repeated the experiment in BL/6 mice that had been treated with an NK1.1-depleting antibody before the first intracranial injection (**Fig. 6A**). IL-15c administration did not restore NK1.1^+^ cells after depletion and induced a less effective anti-metastatic effect in these mice (**Fig. 6H-I**, **Fig. S14A**). In Rag1^-/-^ mice, NK1.1 depletion completely abrogated the anti-tumor effect of IL-15c treatment (**Fig. 6J-K**). Together, these data indicate that NK cells are the principal effector population mediating IL-15c-induced protection, while adaptive immune cells can make a partial additional contribution in immunocompetent mice.

**Figure 6.**
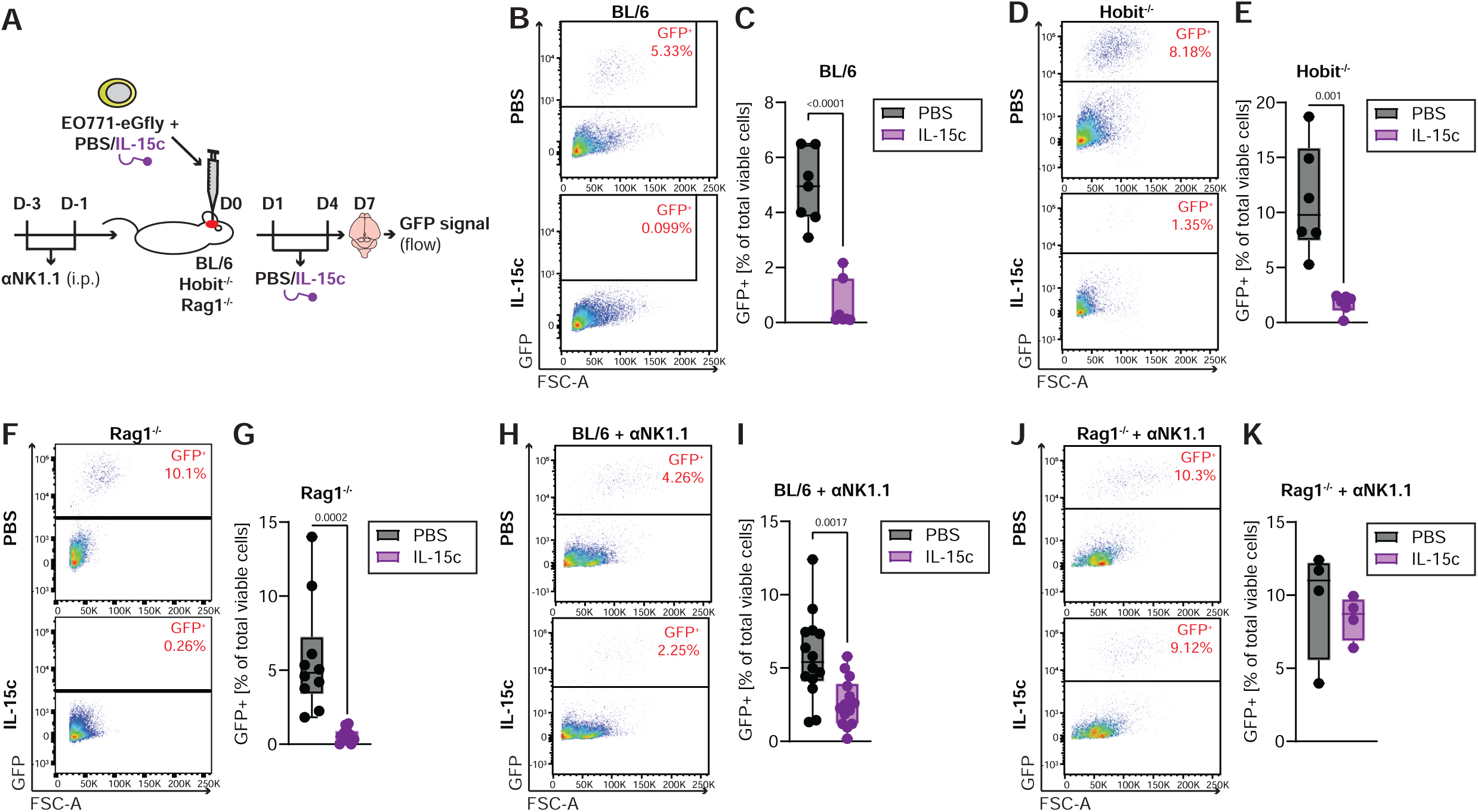
Group 1 ILCs are critical to limit brain metastasis upon IL-15c treatment. **(A)** Experimental schematic for IL-15 agonist treatment. BL/6, *Hobit*^−/−^, or *Rag1*^−/−^ mice received intracranial EO771 eGfly cells together with PBS (control) or IL-15 complex (IL-15c). Mice received additional PBS or IL-15c doses on days 1 and 4 post-tumor implantation and were euthanized on day 7 for flow cytometric assessment of tumor burden. In selected experiments, BL/6 mice (H–I) or *Rag1*^−/−^ mice (J–K) were treated with αNK1.1 antibody 3 and 1 days prior to EO771 eGfly injection to deplete NK1.1^+^ cells. **(B-C)** Flow cytometry gating strategy (B) and quantification (C) of GFP^+^ EO771 eGfly tumor cells in digested brains of BL/6 mice that were either treated with PBS (n=7) or IL-15c (n=7). Cells were pre-gated on viable cells. **(D-E)** Flow cytometry gating strategy (D) and quantification (E) of GFP^+^ EO771 eGfly tumor cells in digested brains of Hobit^-/-^ mice that were either treated with PBS (n=6) or IL-15c (n=6). Cells were pre-gated on viable cells. **(F-G)** Flow cytometry gating strategy (F) and quantification (G) of GFP^+^ EO771 eGfly tumor cells in digested brains of Rag1^-/-^ mice that were either treated with PBS (n=5) or IL-15c (n=5). Cells were pre-gated on viable cells. **(H-I)** Flow cytometry gating strategy (H) and quantification (I) of GFP^+^ EO771 eGfly tumor cells in digested brains of BL/6 mice that were either treated with PBS (n=14) or IL-15c (n=14). Cells were pre-gated on viable cells. BL/6 mice were treated with an αNK1.1 antibody to deplete NK1.1 cells before injection with EO771 eGfly cells. **(J-K)** Flow cytometry gating strategy (J) and quantification (K) of GFP^+^ EO771 eGfly tumor cells in digested brains of Rag1^-/-^ mice that were either treated with PBS (n=4) or IL-15c (n=4). Cells were pre-gated on viable cells. Rag1-/-mice were treated with an αNK1.1 antibody to deplete NK1.1 cells before injecting EO771 eGfly cells. Data are represented as mean ± SD. Statistics: two-tailed Student’s t-test (C, E, G, I). Exact p-values are shown for statistically significant comparisons.

### iPSC-derived NK cell therapy improves control of brain metastasis

Several NK cell therapy approaches have been developed in recent years, including off-the-shelf umbilical cord blood or iPSC-derived NK cell therapies^12,64–67^. These NK cells can be genetically engineered to express and secrete IL-15, an autocrine activator that enhances their activity and anti-tumor efficacy^67^. Since we observed improved brain metastasis control post IL-15c treatment, an effect mediated by NK cells, we aimed to test the potential of iPSC-derived IL-15-secreting NK cells to control brain metastases *in vivo*. Therefore, we intracranially injected the eGFly-tagged human breast cancer cell line MDA-231 into the brain of NSG mice and treated those mice 4 days later with PBS (control group), wild-type (wt) iPSC-derived NK cells, or IL-15-secreting iPSC-derived NK cells (**Fig. 7A**). The PBS and NK cells were injected intracranially. Eleven days after NK cell injection, we observed reduced metastatic burden in mice injected with IL-15-secreting NK cells (experiments 1 and 2) and wt NK cells (experiment 2), measured by *ex vivo* BLI and flow cytometry (**Fig. 7B-E**, **Fig. S15A-D**). The NK cells showed a greater anti-metastatic effect in the second experiment. Eleven days post-injection, only the IL-15-secreting NK cells were retained in the brains of the mice, while the wt NK cells were no longer detectable by flow cytometry (**Fig. 7F-G**, **Fig. S15E-F**). The frequency of retained iPSC-derived IL-15-secreting NK cells was higher in the second experiment than in the first, potentially explaining the greater anti-metastatic effect seen in this experiment. Interestingly, wt NK cell-treated mice showed the same reduction in metastatic burden as IL-15-secreting NK cell-treated mice, even though no NK cells were detectable at the endpoint. Potentially, wt NK cells survived long enough in the brain to reduce metastatic burden efficiently but could not survive long-term without IL-15 supplementation. A longer experiment is needed to test whether wt and IL-15-secreting NK cells differ in controlling brain metastasis over time.

**Figure 7.**
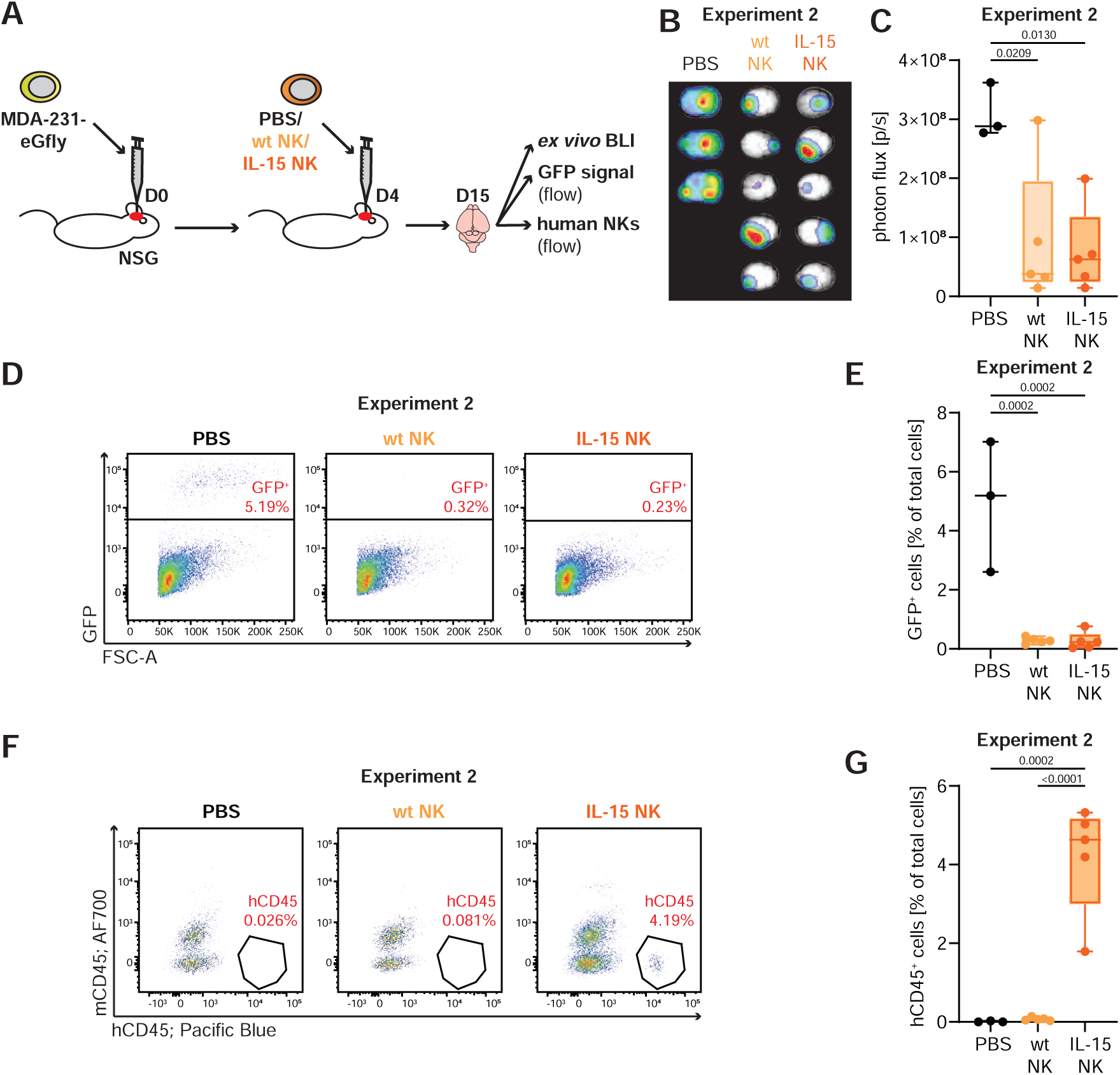
iPSC-derived NK cell therapy improves control of brain metastasis. **(A)** Experimental schematic for treatment of brain metastases with iPSC-derived NK cells. NSG mice received intracranial MDA-MB-231 eGfly cells. Four days later, mice were treated intracranially with PBS (control; n=4 Experiment 1, n=3 Experiment 2), iPSC-derived wild-type NK cells (wt NK; n=5 Experiment 2), or iPSC-derived IL-15-producing NK cells (IL-15 NK; n=4 Experiment 1, n=5 Experiment 2). Mice were euthanized on day 15 post-tumor implantation. Tumor burden was monitored by bioluminescence imaging (BLI) and flow cytometry, and human NK cell abundance was assessed at endpoint. **(B-C)** *Ex vivo* BLI of brains harvested on day 15 post-implantation (B) and corresponding quantification of total bioluminescence signal (C) from PBS-treated controls and iPSC-derived NK cell–treated mice. **(D-E)** Flow cytometry gating strategy (D) and quantification (E) of GFP^+^ MDA-MB-231 eGfly tumor cells in digested brains from PBS-treated controls and iPSC-derived NK cell–treated NSG mice. Cells were pre-gated on viable cells. **(F-G)** Flow cytometry gating strategy (F) and quantification (G) of human NK cells (mCD45^−^hCD45^+^) in brains from PBS-treated and iPSC-derived NK cell–treated NSG mice. Cells were pre-gated on viable cells; data are shown as the frequency of hCD45^+^ cells among all viable cells. Data are represented as mean ± SD. Statistics: one-way ANOVA (C, E, G). Exact p-values are shown for statistically significant comparisons.

## DISCUSSION

In this study, we investigated whether microenvironmental differences shape group 1 ILC responses to metastatic cancer. By combining functional depletion and genetic models with scRNA-seq, we identified striking differences in both group 1 ILC composition and anti-metastatic activity between the brain and liver. Whereas the liver contained multiple NK cell and tissue-resident ILC1 populations, approximately 70% of group 1 ILCs in the brain were assigned to a circulating NK cell population, indicating a substantially more restricted repertoire. This designation is based on transcriptional state rather than anatomical location, so it should not be interpreted as direct evidence that these cells remain within the vasculature. However, our flow cytometry-based intravascular-labeling experiments support the conclusion that most brain-associated NK cells are vascular rather than parenchymal. Functionally, depletion of group 1 ILCs increased metastatic burden in the liver but had little effect on BCBM. This is consistent with the established role of NK cells and liver-resident ILC1s in hepatic metastatic surveillance, where these populations provide complementary control of metastatic seeding and outgrowth^41,42^. In BCBM, NK-cell abundance appears limited and variable across studies, with some datasets reporting markedly reduced or undetectable NK-cell populations compared with primary breast tumors^68,69^. Nevertheless, preclinical studies have demonstrated that engineered NK-cell approaches can exert anti-tumor activity against BCBM models, although whether endogenous NK cells or tissue-resident ILC1s provide physiologically relevant surveillance against BCBM remains undefined^70^. Our findings identify a pronounced site-specific failure of endogenous group 1 ILC-mediated metastatic control in the brain. These conclusions are based on experimental mouse metastasis models and therefore may not capture the full biological and clinical heterogeneity of spontaneous human BCBM.

The limited anti-metastatic activity we observed does not, however, reflect a general inability of group 1 ILCs to respond within the CNS. NK cells and ILC1s occupy distinct anatomical niches in the healthy brain, with ILC1s enriched at CNS border regions such as the choroid plexus and smaller populations present in the meninges and parenchyma^71^. During experimental autoimmune encephalomyelitis, NK cells accumulate and increase IFN-γ and TNF-α production, while ILC1s redistribute toward the inflamed brain parenchyma, demonstrating that both populations can respond dynamically to CNS inflammation^71^. Similarly, NK cells and ILC1s are recruited into the brain during viral infection, although in congenital murine cytomegalovirus infection this response fails to efficiently control viral replication and instead contributes to IFN-γ-dependent neuropathology^72^. A role for group 1 ILCs has also been described in glioblastoma, where infiltrating NK cells show reduced expression of activating receptors and features of TGF-β-mediated suppression despite retaining the capacity to recognize glioblastoma cells *ex vivo*^73,74^. Moreover, in melanoma brain metastasis, NK cells show limited direct cytotoxicity but are required for effective checkpoint blockade and CD8^+^ T cell recruitment^75^. Together with our finding that most brain-associated NK cells remain intravascular, these studies suggest that the CNS can support group 1 ILC recruitment and activation, but that tissue-specific constraints on parenchymal access, persistence, or functional differentiation may limit effective anti-metastatic surveillance.

We therefore hypothesized that differences in tissue-derived signals required for group 1 ILC maintenance and activation may contribute to their impaired anti-metastatic response in the brain. We focused on IL-15 and IL-15Rα, as IL-15 is a critical regulator of NK-cell and ILC1 development, survival, and activation and is predominantly delivered through IL-15Rα-mediated trans-presentation to cells expressing IL-2/15Rβ and the common γ-chain^53^. IL-15 is maintained at very low, primarily intracellular levels in the brain with restricted anatomical and cellular expression largely confined to myeloid cells and microglia^76,77^. Expression robustly spikes during pathological stress, particularly under inflammatory conditions, where the primary source of IL-15 shifts to astrocytes^78^. While this has largely been described in inflammatory states in the brain such as Multiple Sclerosis, it is unknown whether this signaling cascade is deployed in brain malignancies^78,79^. IL-15Rα is required to efficiently stabilize and present bioactive IL-15, and IL-15/IL-15Rα complexes provide substantially more potent and sustained stimulation than soluble IL-15 alone^54^. Consistent with this model, we found markedly higher IL-15Rα expression in the liver than in the brain. The liver provides a particularly supportive IL-15 niche, where Kupffer cells, stellate cells, and hepatocytes contribute to local IL-15 availability, and hepatocyte- and macrophage-specific IL-15Rα expression is required to maintain hepatic NK and NKT-cell populations^80,81^. Interestingly, our single-cell analysis identified IL-15Rα expression in a distinct hepatocyte population, suggesting that non-hematopoietic liver cells may directly support tissue-specific group 1 ILCs. This local IL-15 trans-presentation could help sustain the abundant and diverse group 1 ILC compartment required for immune surveillance in the liver. In contrast, reduced IL-15Rα availability in the brain may limit local IL-15 trans-presentation and thereby restrict the survival, expansion, or functional differentiation of group 1 ILCs after entering the parenchyma. Although the association between reduced brain IL-15Rɑ and impaired group 1 ILC function is strengthened by the therapeutic response to IL-15/IL-15Rɑ complexes, our experiments do not establish a direct causal link between reduced endogenous IL-15Rɑ and failed brain metastasis surveillance. Direct manipulation of the IL-15Rɑ availability in the brain and liver will be required to test this mechanism definitively. Whether this relatively restricted cytokine environment represents an adaptation that limits potentially damaging innate immune activation in the CNS remains speculative, but it is consistent with the need for particularly tight regulation of inflammatory responses in neural tissues.

To overcome the limited IL-15/IL-15Rα signaling in the brain, we administered IL-15/IL-15Rα complexes locally to tumor-bearing mice and found that this treatment restored group 1 ILC expansion and improved control of BCBM. IL-15 has long been investigated as a cancer immunotherapy because it expands and activates NK and CD8^+^ T cells, but early clinical studies of systemic recombinant IL-15 revealed substantial dose- and schedule-dependent toxicities, including fever, hypotension, thrombocytopenia, and liver enzyme elevations^82–85^. More recent studies have therefore focused on stabilized IL-15/IL-15Rα complexes, receptor agonists, altered dosing schedules, and targeted or localized delivery^85–87^. Notably, the IL-15 superagonist N-803 has now received FDA approval in combination with BCG for non-muscle-invasive bladder cancer, where it is administered locally by intravesical instillation, demonstrating the clinical feasibility of compartment-restricted IL-15-based immunotherapy^88^. Local IL-15 delivery may be particularly attractive for CNS tumors, where systemic exposure is undesirable, and BBB penetration can be limiting. Indeed, direct cerebral IL-15 infusion, IL-15-producing microglia, and locally delivered IL-15/IL-15Rα-expressing oncolytic viruses have each enhanced antitumor immunity in preclinical glioma models^58,89–91^. In our study, local IL-15-complex treatment produced no overt neurological toxicity during the seven-day treatment period. However, longer-term studies incorporating dedicated neuropathological and systemic safety assessments will be required before the safety of repeated CNS administration can be established. Additionally, direct intracranial injection is useful for establishing biological proof of concept but is clinically invasive. It is crucial to develop delivery strategies that achieve sufficient CNS exposure while remaining feasible for repeated administration in patients. Future studies should also determine the dosing frequency required to maintain metastatic control and whether IL-15-complex therapy can be combined with established treatments such as radiotherapy or immune checkpoint blockade.

To further translate our findings, we evaluated the potential of human iPSC-derived NK cells engineered to constitutively secrete IL-15 to control BCBM. A single intracranial administration significantly reduced BCBM burden, and IL-15-secreting iPSC-NK cells remained detectable in the brain 11 days after treatment. iPSC-derived NK cells are an attractive platform for adoptive immunotherapy because they can be generated as standardized, scalable, off-the-shelf products and genetically engineered to enhance tumor recognition, persistence, and resistance to immunosuppression^64,66^. Preclinical studies have demonstrated activity of iPSC-NK cells against both hematological malignancies and solid tumors, including B-cell lymphoma and ovarian cancer, and several engineered iPSC-NK products have subsequently entered early-phase clinical testing^64,66^. Notably, recent work with the clinical-grade iPSC-derived NK-cell product FT538 demonstrated intracranial persistence and anti-tumor activity in orthotopic glioblastoma models, further supporting this platform’s feasibility for CNS malignancies^92^. To our knowledge, however, iPSC-derived NK cells have not previously been evaluated in BCBM, making our findings the first evidence that these cells can exert therapeutic activity and persist within the BCBM microenvironment. Several limitations remain. Our experiments were restricted to an 11-day observation period and a single intracranial NK-cell administration, and therefore do not establish the duration of metastatic control, optimal cell dose or dosing frequency, or long-term persistence and safety. Furthermore, non-IL-15-secreting iPSC-NK cells also reduced metastatic burden despite being undetectable at endpoint, suggesting that transient NK-cell activity may be sufficient to eliminate a proportion of tumor cells early after treatment. Longer-term studies directly comparing IL-15-secreting and unmodified iPSC-NK cells, including repeated dosing and longitudinal assessment of NK-cell persistence and tumor recurrence, will therefore be required to determine whether autonomous IL-15 production provides a durable therapeutic advantage

In summary, our study identifies a tissue-specific failure of group 1 ILC-mediated metastatic surveillance in the brain, supporting a model in which limited IL-15/IL-15Rα signaling contributes to this impaired control. In contrast, the liver supports a more diverse and functionally effective group 1 ILC compartment that restricts metastatic outgrowth. Importantly, this site-specific defect was therapeutically reversible. Local delivery of IL-15/IL-15Rα complexes restored anti-metastatic activity, while IL-15-secreting iPSC-derived NK cells provided an alternative strategy to deliver sustained NK-cell stimulation directly within the brain. Together, these findings establish the IL-15/IL-15Rα axis as a potential therapeutic vulnerability in BCBM and provide a rationale for developing localized cytokine-based or engineered NK-cell approaches to restore innate immune surveillance in the CNS. Further studies are needed to define optimal dosing, durability, safety, and combinatorial treatment strategies before translating these approaches to clinical evaluation in patients with BCBM.

## MATERIALS AND METHODS

### Cell lines

EO771 (ATCC CRL-3461) and MDA-MB-231 (ATCC HTB-26) cells were obtained from the American Type Culture Collection (ATCC). The pCDH–EF1α–eFFly–eGFP plasmid (Addgene plasmid #104834; RRID) was kindly provided by Irmela Jeremias. To generate lentiviral particles, we co-transfected HEK293T cells with pCDH–EF1α–eFFly–eGFP and the packaging plasmids pMD2.G and psPAX2 using Lipofectamine 2000 (Invitrogen, cat. no. 11668027). We collected viral supernatants and used them to transduce EO771 and MDA-MB-231 cells overnight with 8 μg/mL polybrene (Sigma-Aldrich, cat. no. TR-1003-G). Successfully transduced cells were isolated based on GFP expression using a BD FACSAria Fusion cell sorter and are hereafter referred to as EO771-eGFly and MDA-MB-231-eGFly.

EO771-eGFly cells were cultured in RPMI 1640 supplemented with 5% (vol/vol) FBS, 10 U/mL penicillin, 0.1 mg/mL streptomycin, and 10 mM HEPES at 37 °C in a humidified incubator containing 5% CO₂. MDA-MB-231-eGFly cells were cultured in DMEM supplemented with 10% (vol/vol) FBS, 10 U/mL penicillin, and 0.1 mg/mL streptomycin under the same incubation conditions. The identity of the tumor cell lines was confirmed by short tandem repeat (STR) profiling by ATCC before use in animal experiments.

### Mice

C57BL/6J (BL/6; RRID:IMSR_JAX:000664), B6(Cg)-Tyrc-2J/J (BL/6 albino; RRID:IMSR_JAX:000058), B6.129S7-Rag1^tm1Mom^/J (Rag1^-/-^; RRID:IMSR_JAX:002216), and NOD.Cg-Prkdc^scid^ Il2rg^tm1Wjl/Sz^J (NSG; RRID:IMSR_JAX:005557) mice were obtained from The Jackson Laboratory. Hobit^-/-^ mice were kindly provided by Timothy E. O’Sullivan (University of California, Los Angeles).

Mice were housed in individually ventilated cages in an enriched environment under controlled temperature and humidity in a vivarium overseen by University Laboratory Animal Resources (ULAR) at the University of California, Irvine. ULAR personnel provided veterinary care. We used female mice for all experiments since the overwhelming majority of breast cancers arise in women. Mice between the ages of 4-6 weeks were used for the intracranial injections; for the other experiments, mice between the ages of 5-8 weeks were included in the experiments. The Institutional Animal Care and Use Committee (IACUC) at the University of California, Irvine, reviewed and approved all animal procedures, which were performed under UCI IACUC protocols 19-051, 22-033, and 25-015.

### Generation of iPSC-derived NK cells

Human induced pluripotent stem cells (iPSCs) were genetically engineered to express human IL-15 and differentiated toward the NK-cell lineage using a previously described protocol with minor modifications^93^. During hematopoietic induction and NK-cell differentiation, cells were cultured with stem cell factor (SCF), FLT3 ligand (FLT3L), IL-7, and IL-15. Following NK-cell differentiation, iPSC-derived NK (iPSC-NK) cells were further expanded by co-culture with irradiated K562-based artificial antigen-presenting cells (aAPCs) in the presence of IL-2. SCF and FLT3L were obtained from Innocyto, whereas IL-2, IL-7, and IL-15 were obtained from the NIH Biological Resources Branch. The iPSC-NK cells were characterized for NK-cell phenotype prior to downstream experiments.

### Human specimens

Archival formalin-fixed paraffin-embedded human tissue was obtained under a protocol approved by the University of California, Irvine Institutional Review Board (UCI 17-05). Samples comprised non-neoplastic brain (epilepsy resection, n=10), breast-cancer brain metastasis (n=26), non-neoplastic liver (n=10), and liver metastasis (n=11). Section thickness was 4–5 µm for each slide analyzed.

### Antibody-mediated depletion of NK1.1^+^ cells

To deplete NK1.1^+^ cells, mice received 200 μg anti-NK1.1 antibody (clone PK136; BioLegend, cat. no. 108761) by intraperitoneal injection on days -3 and -1 relative to tumor cell injection. Where indicated, mice received an additional 200 μg dose on day 4 after tumor cell injection. Control mice received an equivalent volume of 1x PBS. We confirmed NK1.1^+^ cell depletion at the experimental endpoint by flow cytometry.

### Intracardiac tumor cell injection

Intracardiac injections were performed as described by Campbell et al.^94^ into the left cardiac ventricle of mice. Mice were anesthetized with 2% isoflurane in 100% oxygen, and 0.1mg carprofen was administered intraperitoneally for analgesia. Tumor cell injections were performed under ultrasound guidance using a Butterfly iQ ultrasound device (Butterfly Network, Inc.) and sterile ultrasound gel (Parker Labs). EO771-eGFly cells were resuspended in chilled PBS, and 5 × 10^5^ cells in a total volume of 100 μL were loaded into a 1-mL syringe equipped with a 28-G, ½-inch needle. The needle was visualized in-plane by ultrasound and advanced into the left ventricular cavity. After confirmation of correct needle placement, the cell suspension was injected, and the needle was withdrawn. Mice were transferred to a heated recovery cage and monitored until recovery from anesthesia.

### Intracranial cell injection

EO771-eGFly cells were resuspended at 10 × 10^6^ cells/mL in chilled PBS. MDA-MB-231-eGFly cells were resuspended at 5 × 10^6^ cells/mL in chilled PBS. Mice were anesthetized with 2% isoflurane in 100% oxygen, and 0.1mg carprofen was administered intraperitoneally for analgesia. The head was sterilized, and intracranial injections were performed using a 10 μL glass Hamilton syringe equipped with a removable 31-G needle (Hamilton Company). The syringe was rinsed with ethanol, then PBS, before use. A total volume of 10 μL containing 1 × 10^5^ EO771-eGFly cells or 0.5 x 10^5^ MDA-MB-231-eGFly cells was injected laterally through the patent coronal suture of 4–6-week-old mice. The needle was advanced to a depth of 3 mm using a plastic spacer as a guide, and the plunger was depressed over approximately 20 s. After injection, we withdrew the needle, disinfected the injection site, and transferred mice to a heated recovery cage, where we monitored them until recovery from anesthesia.

Four days post-intracranial injection of the MDA-MB-231-eGFly cells into the NSG mice, 5 x 10^6^ human iPSC-derived NK cells were injected intracranially into the same site, following the same procedure as described above. For this experiment, mice were divided into three groups and injected with iPSC-derived wild-type NK cells, iPSC-derived IL-15-secreting NK cells, or PBS. Fifteen days post-tumor cell injection, mice were sacrificed, and the presence of the iPSC-derived NK cells was tested by flow cytometry.

### IL-15 agonist preparation and treatment

IL-15 complexes (IL-15c) were generated using recombinant murine IL-15 (PeproTech/Thermo Fisher Scientific, cat. no. 210-15) and recombinant murine IL-15Rα-Fc (R&D Systems, cat. no. 551-MR). Recombinant IL-15 was reconstituted and stored as aliquots containing 2 μg IL-15 in 20 μL PBS, whereas IL-15Rα-Fc was stored as aliquots containing 10 μg IL-15Rα-Fc in 20 μL PBS. Equal volumes of the two solutions were combined, resulting in an IL15:IL-15α-Fc mass ratio of 1:5, and incubated at 37 °C for 30 min to allow complex formation.

For intracranial co-injections with tumor cells, 100 µL of EO771-eGFly cells resuspended at 10^6^ cells/mL were combined with 40 µL IL-15c. For the control mice, the EO771-eGFly suspension was supplemented with 40 µL 1x PBS to maintain identical EO771-eGFly cell concentrations between the control and IL-15c groups. We injected 10 µL of each mix intracranially. For intracranial doses of IL-15c administered without cells, 40 µL IL-15c was diluted in 100 µL 1x PBS and subsequently administered to each mouse as a 10 µL injection. All injections were performed as described in the paragraph ‘*intracranial cell injection*’, with the IL-15c/PBS injections done on the same site as the EO771-eGFly/IL-15c co-injections.

### Bioluminescence imaging

We monitored *in vivo* tumor burden longitudinally by bioluminescence imaging (BLI) at the indicated experimental time points. When necessary, we removed fur overlying the imaging region using an electric shaver. Mice were anesthetized with isoflurane and administered D-luciferin (150 mg/kg; GoldBio) intraperitoneally in sterile PBS. Ten minutes after D-luciferin administration, mice were imaged using an IVIS Lumina III In Vivo Imaging System (Xenogen) or an AMI-HT imaging system (Spectral Instruments Imaging).

Regions of interest (ROIs) were defined over the head and body, as appropriate for the experiment. Bioluminescence was quantified as average radiance (photons s^-1 cm^-2 sr^-1) using Living Image software (Xenogen) or Aura software (Spectral Instruments Imaging).

For *ex vivo* imaging, we collected brains and livers at the experimental endpoint and incubated them for 10 min in chilled PBS containing 2% FBS and 0.3 mg/mL D-luciferin (GoldBio, cat. no. LUCK-1G).

Organs were subsequently imaged. We defined ROIs encompassing the respective organs and quantified average radiance as described above.

### Intravascular CD45 labeling

To distinguish intravascular from tissue-localized immune cells, mice received anti-CD45 conjugated to PE-Cy5 (clone 30-F11, Biolegend cat. #103110) intravenously immediately before euthanasia. The antibody was diluted to 2 μg/mL in sterile PBS, and 100 μL was administered through the tail vein, corresponding to 0.2 μg antibody per mouse. Mice were warmed for approximately 20 min before injection to facilitate tail vein visualization. After intravenous administration, we allowed the antibody to circulate for 3 min before euthanizing mice by CO₂ asphyxiation. During subsequent *ex vivo* surface staining, we used a second anti-CD45 antibody conjugated to Alexa Fluor 700 to identify total CD45^+^ leukocytes. Intravascular cells were therefore identified as PE-Cy5^+^ among total CD45-AF700^+^ leukocytes, whereas PE-Cy5-negative CD45-AF700^+^ cells were considered extravascular at the time of antibody administration. We analyzed samples using a BD LSRFortessa X-20 flow cytometer.

### Tissue digestion

Before tissue collection, mice were transcardially perfused with 20 mL 0.01 M EDTA in 1x PBS.

Whole brains were dissected, sectioned, and transferred to gentleMACS C Tubes (Miltenyi Biotec, cat. no. 130-093-237) containing HBSS++ supplemented with 1 mg/mL Collagenase D (MilliporeSigma, cat. no. 11213857001) and 10 μg/mL DNase I (Fisher Scientific, cat. no. NC9709009). Brain tissue was dissociated using a gentleMACS Octo Dissociator with Heaters according to the 37 °C adult brain dissociation program (ABDK1-37C). Following dissociation, cell suspensions were filtered through 70-μm cell strainers and centrifuged at 300 × g for 5 min at 4 °C. Myelin and cellular debris were removed using Debris Removal Solution according to the manufacturer’s instructions (Miltenyi Biotec, cat. no. 130-109-398). Residual erythrocytes were lysed using ACK Lysis Buffer (Gibco, cat. no. A10492-01). Cells were subsequently counted and used for downstream applications.

The left lobe of the liver was mechanically minced with a razor blade in 0.5 mL digestion buffer containing 1mg/mL Collagenase D and 10μg/mL DNase1 in 1x HBSS. Tissue homogenates were transferred to 15-mL conical tubes, then digestion buffer was added to a final volume of 7 mL. Samples were incubated at 37 °C for 20 min with continuous shaking at 200 rpm. Following digestion, cell suspensions were filtered through 70-μm cell strainers. Cellular debris was removed using Debris Removal Solution according to the manufacturer’s instructions (Miltenyi Biotec, cat. no. 130-109-398). Erythrocytes were lysed using ACK Lysis Buffer, after which cells were counted and used for downstream applications.

### Flow cytometric analysis of group 1 ILCs and GFP^+^ tumor cells

Single-cell suspensions generated from brain and liver tissues were analyzed by flow cytometry. A maximum of 1 × 10^6^ cells per sample was stained in 50 μL FACS buffer consisting of PBS supplemented with 2% (vol/vol) FBS.

To analyze group 1 ILCs, cells were incubated with anti-CD16/32 (1:50; BioLegend, cat. no. 101302) for 10 min at 4 °C to block Fc receptors. Cells were subsequently incubated for 30 min at 4 °C with a biotinylated lineage antibody cocktail containing antibodies against CD19 (1:150; BioLegend, cat. no. 115504), Gr-1 (1:150; BioLegend, cat. no. 108404), TER119 (1:150; BioLegend, cat. no. 116204), EpCAM (1:150; BioLegend, cat. no. 118204), CD4 (1:150; BioLegend, cat. no. 100404), and CD8 (1:150; BioLegend, cat. no. 100704). Cells were washed with FACS buffer and centrifuged at 300 × g for 5 min at 4 °C. Cells were then stained for 30 min at 4 °C with Live/Dead Fixable Aqua (1:100; Thermo Fisher Scientific, cat. no. L34957), CD45-Alexa Fluor 700 (1:100; BioLegend, cat. no. 103128), NK1.1-APC (1:100; Invitrogen, cat. no. 17-5941-81), NK1.1-APC-Cy7 (1:100; BioLegend, cat. no. 108126), NK1.1-PE-Cy7 (1:100; BioLegend cat. no. 108714), CD49b-PE-Texas Red (1:100; BioLegend, cat. no. 108924), CD49a-BUV395 (1:100; BD Biosciences, cat. no. 740262), and streptavidin-BV650 (1:1,000; BioLegend, cat. no. 405232). Cells were subsequently washed and resuspended in FACS buffer for acquisition.

For flow cytometric validation of human iPSC-derived NK cell retention, we stained the brain single-cell suspensions with anti-CD16/32 (1:50; BioLegend, cat. no. 101302) for 10 min at 4 °C to block Fc receptors. Cells were subsequently incubated for 30 min at 4 °C with a cell surface antibody mix, containing Live/Dead Fixable Aqua (1:50; Thermo Fisher Scientific, cat. no. L34957), mouse CD45-Alexa Fluor 700 (1:50; BioLegend, cat. no. 103128), human CD45-Pacific Blue (1:50; BioLegend, cat. no. 304021), and human CD56-BV650 (1:50; BioLegend, cat. no. 362531). Cells were subsequently washed and resuspended in FACS buffer for acquisition.

For flow cytometric quantification of GFP^+^ tumor cells, 30% of each brain or liver single-cell suspension was allocated to tumor-burden analysis. Cells were stained with Live/Dead Fixable Aqua (Thermo Fisher Scientific, cat. no. L34957), washed, and resuspended in 300 μL chilled FACS buffer. We determined the frequency of viable GFP^+^ cells by flow cytometry and used it as a measure of tumor burden.

We acquired samples on a BD LSRFortessa X-20 flow cytometer using FACSDiva software (BD Biosciences) and analyzed them in FlowJo v10.10. We included appropriate fluorescence-minus-one controls where required for gate definition.

### Lineage depletion by magnetic-activated cell sorting

To enrich group 1 ILCs from brain and liver single-cell suspensions, lineage-positive cells were depleted by magnetic-activated cell sorting (MACS). All staining and washing steps were performed in MACS buffer consisting of PBS supplemented with 5% (vol/vol) FBS.

Cells were first incubated with anti-CD16/32 (1:50; BioLegend, cat. no. 101302) for 10 min at 4 °C. Cells were subsequently stained for 30 min at 4 °C with a biotinylated lineage antibody cocktail containing antibodies against CD19 (1:150; BioLegend, cat. no. 115504), Gr-1 (1:150; BioLegend, cat. no. 108404), TER119 (1:150; BioLegend, cat. no. 116204), EpCAM (1:150; BioLegend, cat. no. 118204), CD4 (1:150; BioLegend, cat. no. 100404), and CD8 (1:150; BioLegend, cat. no. 100704), using a final staining volume of 25 μL per 1 × 10^6^ cells.

Following washing and centrifugation at 300 × g for 5 min at 4 °C, cells were resuspended in MACS buffer containing anti-biotin MicroBeads (Miltenyi Biotec, cat. no. 130-090-485) at a 1:4 dilution, using 6.25 μL total staining solution per 1 × 10^6^ cells. After 15 min incubation at 4 °C, cells were washed and resuspended in 0.5–1 mL MACS buffer to a maximum concentration of 1 × 10^8^ cells/mL.

Lineage-positive cells were depleted using either an autoMACS cell separator (Miltenyi Biotec) or LS Columns (Miltenyi Biotec, cat. no. 130-042-401) mounted on a MidiMACS separator (Miltenyi Biotec, cat. no. 130-042-302). We collected the lineage-negative fraction and used it for downstream applications.

### Western blot

Brains and the left liver lobes from healthy BL/6 animals and BL/6 mice 9 days post intracardiac EO771-eGFly injection were harvested and instantly snap-frozen in liquid nitrogen. The collected tissues were then mechanically dissociated using a tissue homogenizer in RIPA buffer (Thermo Fisher, 89900) containing protease and phosphatase inhibitors (Thermo Fisher, 78430) (100 ul / 30 mg of liver tissue, 100 ul / 50 mg of brain tissue). Lysates were incubated with agitation for 45 minutes, followed by centrifugation at 13,000g for 15 minutes. Supernatant was removed, and protein was quantified using Pierce BCA Protein Assay Kit (Thermo Fisher, 23225) according to the manufacturer’s instructions. 75 μg of protein samples were run on a 12% SDS-PAGE gel (BioRad, 4568046), and proteins were wet-transferred to a PVDF membrane. Membranes were blocked with 5% BSA diluted in TBS containing 0.1% Tween-20 (5% BSA-TBST) for one hour at room temperature. Primary antibodies diluted in 5% BSA-TBST were added to the membranes and incubated at 4°C overnight. Primary antibodies included: 1:2000 goat anti-mouse IL-15rɑ (R&D systems, AF551) and 1:1000 mouse anti-mouse Vinculin (Millipore Sigma, V9131). The following day, membranes were washed in TBS containing 0.1% Tween-20 (TBST) followed by incubation with secondary antibodies diluted in 5% BSA-TBST for one hour at room temperature. Secondary antibodies included: 1:2000 Rabbit Anti-Goat HRP (Invitrogen, Cat No. 811620) or 1:1000 Goat Anti-Mouse HRP (Thermo Fisher, Cat No. 31430). Membranes were washed again with TBST and imaged. Protein bands were visualized using Thermo Fisher Chemiluminescent Substrate kit (Thermo Fisher, Cat No. 34579) and imaged on a BioRad ChemiDoc system.

### Generation of group 1 ILC-enriched murine brain and liver scRNA-seq datasets

We generated two group 1 ILC-enriched single-cell RNA-sequencing datasets. For the TimeCourse dataset, BL/6 mice received intracardiac injections of EO771-eGFly cells as described above, whereas control mice received PBS. Mice were euthanized on days 4 and 10 after injection. At both time points, 3 metastasis-burdened and 2 control mice were sacrificed, and the brains and livers were harvested. For the IL15 dataset, BL/6 mice received intracranial injections of EO771-eGFly cells in combination with IL-15c or PBS on day 0 (n = 5 mice per group), followed by an additional intracranial administration of IL-15c or PBS on day 1. Mice were euthanized on day 4, and brains were collected.

We dissociated tissues and depleted lineage-positive cells by MACS as described above. Lineage-negative cells were counted using a Countess II automated cell counter (Thermo Fisher Scientific) and prepared for Cell Multiplexing Oligo (CMO) labeling and fluorescence-activated cell sorting according to the 10x Genomics Cell Multiplexing Oligo Labeling protocol. We performed all subsequent staining steps in PBS supplemented with 10% (vol/vol) FBS. Between 0.2 × 10^6^ and 2 × 10^6^ cells per sample were incubated with anti-CD16/32 (1:50; BioLegend, cat. no. 101302) for 10 min at 4 °C, followed by staining for 30 min at 4 °C with Live/Dead Fixable Aqua (1:50; Thermo Fisher Scientific, cat. no. L34957), CD45-Alexa Fluor 700 (1:50; BioLegend, cat. no. 103128), NK1.1-PE-Cy7 (1:50; BioLegend, cat. no. 108714), and streptavidin-BV650 (1:500; BioLegend, cat. no. 405232). Cells were washed and resuspended in 100 μL of an assigned CMO from the 3′ CellPlex Kit Set A (10x Genomics, cat. no. 1000261). We labeled each biological sample with a distinct CMO. Following a 5-min incubation at room temperature, cells were washed, centrifuged at 300 × g for 5 min at 4 °C, and resuspended in 280 μL PBS containing 10% FBS for sorting. Viable lineage-negative CD45^+^NK1.1^+^ cells were sorted using a BD FACSAria II cell sorter into chilled PBS containing 10% FBS. Biological samples were sorted separately but collected into a common tube after CMO labeling to generate the respective multiplexed library. Sorted cells were centrifuged and resuspended at the appropriate concentration for loading onto the 10x Genomics Chromium platform.

Single-cell libraries were generated according to the Chromium Next GEM Single Cell 3′ Gene Expression user guide (10x Genomics, CG000183 Rev C) using v3.1. We loaded cells to achieve a target recovery of approximately 10,000 cells per library. We generated three libraries: TimeCourse day 4, TimeCourse day 10, and IL15. We sequenced the libraries on an Illumina NovaSeq 6000 to a target depth of approximately 50,000 reads per cell for the gene-expression libraries. We processed sequencing reads with Cell Ranger (10x Genomics) v6.1.2 for the TimeCourse dataset and v7.0.1 for the IL15 dataset^95^. Samples within each library were multiplexed with 3′ CellPlex Cell Multiplexing Oligos (CMO301– CMO312), so we aligned and demultiplexed reads to their source sample using cellranger multi against the mm10 (GRCm38) 2020-A mouse reference (refdata-gex-mm10-2020-A for the TimeCourse dataset; refdata-cellranger-arc-mm10-2020-A-2.0.0 for the IL15 dataset). The two TimeCourse libraries were subsequently combined with cellranger aggr.

### Single-cell integration, clustering, and annotation

We combined two independently generated Seurat objects for analysis: the TimeCourse experiment (brain and liver; PBS vs. tumor) and the IL15 experiment (brain only; PBS vs. IL-15c). Each experiment’s cells had already been quality-filtered and doublet-depleted (DoubletFinder) before combination, and each was tagged with its experiment of origin. We merged the two objects into a single object (Seurat v5; counts held as per-experiment layers). We normalized the merged data by log-normalization (NormalizeData, LogNormalize) and identified the 2,000 most variable genes using the variance-stabilizing transform (FindVariableFeatures, vst). We scaled the data (ScaleData) and reduced dimensionality by principal component analysis (RunPCA). Batch effects between the two experiments were then corrected by canonical-correlation-analysis integration using the Seurat v5 layer-based workflow (IntegrateLayers, method = CCAIntegration)^96,97^. Layers were rejoined afterward (JoinLayers). All neighbor-graph, clustering, and embedding steps used the integrated CCA reduction and its first 30 dimensions.

A shared nearest-neighbor graph was built on the integrated reduction (FindNeighbors, dims 1:30), cells were clustered by the Louvain algorithm^98^ (FindClusters, resolution 0.52), and a UMAP embedding was computed on the same 30 integrated dimensions (RunUMAP)^99^. This produced the All-Cells object, which contains both experiments and still includes contaminating T cells and microglia. T-cell clusters and the microglial cluster were then removed, and the retained NK/ILC1 cells were re-clustered on the same integrated CCA embedding (resolution 0.45; UMAP re-computed on the same 30 integrated dimensions). This yielded nine clusters (0–8) of NK and ILC1 cells and defines the root embedding used for all single-cell figures. We annotated clusters manually using canonical NK and ILC1 marker genes.

All single-cell figure panels are subsets of this one root embedding and are not re-clustered or re-integrated. Figure 1 is the root object subset to TimeCourse cells; Figure 5 is the root object with a treatment label (PBS / IL-15c / Liver); brain-specific analyses subset the root object to brain cells without re-clustering. Because every panel shares the same coordinates and cluster identities, cluster colors and positions are directly comparable across figures. We assessed differential expression with the Wilcoxon rank-sum test (FindAllMarkers/FindMarkers) with Bonferroni correction.

### Analysis of publicly available mouse single-cell transcriptomic datasets

To investigate the expression of chemokines involved in NK-cell recruitment and cytokines associated with NK-cell activation, survival, and maintenance under homeostatic conditions, we analyzed publicly available mouse liver and brain single-cell transcriptomic datasets. We evaluated expression of selected genes across annotated cell populations to identify potential tissue-specific cellular sources of NK-cell-regulating factors.

For the liver, publicly available steady-state mouse single-cell RNA-sequencing (scRNA-seq) data were obtained from the Liver Cell Atlas generated by Guilliams et al. (GEO accession GSE192742)^100^. The Mouse Steady State (StSt) dataset comprised a gene-by-cell count matrix and an associated cell annotation matrix containing quality-controlled cells, precomputed UMAP coordinates, cluster identities, and cell-type annotations. The “ALL LIVER CELLS” dataset was used and gene expression was analyzed across the annotated liver cell populations.

For the brain immune compartment, publicly available scRNA-seq data from wild-type homeostatic mouse brains were obtained from the Brain Immune Atlas generated by Van Hove et al. (GEO accession GSE128855)^101^. We used the full aggregate dataset, which includes dura mater, subdural meninges, and choroid plexus samples. The downloaded dataset comprised a gene-by-cell count matrix and corresponding cell annotations, including precomputed dimensionality-reduction coordinates and annotated immune-cell clusters.

Across all datasets, analyses focused on genes encoding factors implicated in NK-cell recruitment and activation or maintenance, including *Cxcl9*, *Cxcl10*, *Ccl3*, *Ccl4*, *Ccl5*, *Cx3cl1*, *Ccl19*, *Ccl21a*, *Cxcl12*, *Il15*, *Il15ra*, *Il12a*, *Il18*, *Il1b*, *Il33*, *Il27*, *Ifnb1*, *Ifng*, *Tgfb1*, *H60b*. We used Seurat (v5.0.3), and summarized expression as dot plots or feature plots within the annotated cell populations. The original cell-type annotations provided by the respective datasets were retained unless otherwise indicated. No re-clustering of the published datasets was performed.

### Immunohistochemistry (IHC) staining of human brain and liver tissues

Human brain and liver formalin-fixed paraffin-embedded sections were melted at 65°C for 1 h, dewaxed in Histo-Clear (National Diagnostics, HS-200), and rehydrated through graded ethanol to water. Heat-induced epitope retrieval was performed in 10 mM sodium citrate buffer (Sigma-Aldrich, C9999) at 95°C for 20 min. Endogenous enzyme activity was quenched with BLOXALL solution (10 min), and sections were blocked in 2.5% normal horse serum (20 min, room temperature). Sections were incubated overnight at 4°C with mouse anti-IL-15 (Invitrogen, MA5-66100, 1:100) and rabbit anti-IL-15Rα (Proteintech, 16744-1-AP, 1:100). Detection used the ImmPRESS Duet Double Staining Polymer Kit (HRP anti-mouse IgG/AP anti-rabbit IgG; Vector Laboratories, MP-7724): sections were incubated with ImmPRESS Duet reagent for 10 min, developed with ImmPACT DAB EqV for 50 s to visualize IL-15 (brown), and then with ImmPACT Vector Red for 3 min to visualize IL-15Rα (magenta), with PBS washes between steps. Sections were counterstained with Mayer’s hematoxylin (10 min), dehydrated, and mounted in Cytoseal XYL. Development times were fixed across the cohort and determined by visual monitoring. Controls processed identically comprised single-stain sections (hematoxylin, DAB or Vector Red alone), used to measure stain vectors for color deconvolution, and secondary-only sections of brain and liver in which primary antibodies were omitted and neither chromogen developed.

### Analysis of IL-15/IL-15Rα stained human brain and liver tissues

Stained sections were digitized on a Keyence slide scanner at 10x (0.75488 µm per pixel) and processed by an automated macro in Fiji/ImageJ (ImageJ 1.54p, ImageJ2 2.16.0) (**Fig. S7A**). Each image was deconvolved into its three chromogens with Colour Deconvolution2 (G. Landini), using background measured per image from unstained glass, OD = −log10(I/I₀), and stain vectors measured on the single-stain control sections, selected on the condition number of the unmixing matrix, which governs noise amplification on inversion (**Fig. S7B**). Tissue was segmented from the original RGB image: the summed optical densities of its red, green, and blue channels gave a stain-agnostic map, thresholded at OD ≥ 0.10, with a chromaticity criterion (≥ 7/255 levels between the strongest and weakest channel) to reject dim glass, followed by morphological opening, removal of components < 0.005 mm², hole filling, and 100 µm erosion to exclude section-margin artifact. Within that eroded mask, each section was sampled using a systematic uniform-random grid of 500 µm square fields at 1250 µm pitch, seeded per image; a field was retained only if ≥ 99.9% of it lay inside. Eight small liver-metastasis specimens were sampled with adapted geometry: seven needle cores with 250 µm fields at 312 µm pitch and 20 µm erosion, and one punch biopsy with 500 µm fields at 625 µm pitch and 40 µm erosion (**Fig. S7C**). Fields were screened for segmentation failure, fragmentation, tears, folds, and dried or detached tissue. Fields were excluded when the largest near-white region exceeded 2% of the field, the very-dark fraction exceeded 1%, or tissue fraction fell below 80% (**Fig. S7D**). Of the 57 tissue sections included, 3,399 fields were analyzed, with a median of 57 fields per section (3 fields minimum per section). Masked pixels were scored IL-15Rα-positive at Vector Red OD ≥ 0.10, a threshold fixed by scanning OD 0.02–0.40 across 21 sections, and IL-15-positive at DAB OD ≥ 0.30. (**Fig. S7E**).

### Statistical Analysis

No statistical methods were used to predetermine sample sizes. Sample sizes were selected based on those commonly used in previous studies employing similar experimental approaches. Unless otherwise indicated, comparisons between two independent groups were performed using unpaired two-tailed Student’s *t*-tests, whereas comparisons involving multiple groups or experimental factors were performed using analysis of variance (ANOVA) followed by appropriate multiple-comparisons tests where applicable. Statistical significance was defined as *P* < 0.05. For flow cytometric analyses, samples with insufficient numbers of viable cells for reliable quantification were excluded from the respective analysis. Where feasible, mice were randomly assigned to experimental groups before initiation of the experiment. Investigators were not blinded to experimental group allocation or outcome assessment unless otherwise indicated. All statistical analyses were performed using GraphPad Prism 10 (GraphPad Software, LLC). Sample sizes, statistical tests, and multiple-comparisons procedures used for individual experiments are provided in the corresponding figure legends.

Analyses of the IL-15/IL-15Rα stained human samples used Python 3.13.12 with SciPy 1.17.1 and NumPy 2.4.3. Because values span several orders of magnitude and are strongly right-skewed, non-parametric tests were used throughout: Kruskal–Wallis across the four groups, followed by two-sided Mann–Whitney *U* tests for all six pairwise comparisons with Benjamini–Hochberg false-discovery-rate correction; adjusted *P* values are reported. No statistical method was used to predetermine sample size, and no randomization or blinding was applicable to this retrospective analysis of archival material.

## Supplementary Materials

**Supplemental information**: containing Supplemental Figures 1-15.

**Supplemental Table 1 (related to Fig. S1):** Marker genes for the NK, NKT, and ILC1 cluster in the ‘TimeCourse’ scRNA-seq dataset.

**Supplementary Table 2 (related to Fig. 1):** Marker genes for the clusters defined in Figure 1 in the ‘TimeCourse’ scRNA-seq dataset.

**Supplementary Table 3 (related to Fig. S3):** List of genes for the Tissue_residency, Circulatory, NFkB-Core, and Stress-IEG module scores.

**Supplementary Table 4 (related to Fig. S3):** Average gene expression for the genes listed in the Tissue_residency, Circulatory, NFkB_Core, and Stress_IEG module scores, split by clusters defined in Figure 1.

**Supplementary Table 5 (related to Fig. S3):** Average module gene expression scores for the Tissue_residency, Circulatory, NFkB_Core, and Stress_IEG modules, split by clusters defined in Figure 1.

**Supplementary Table 6 (related to Fig. 4):** Metadata for the human liver and brain samples.

**Supplementary Table 7 (related to Fig. S10):** Marker genes for the NK, NKT, and ILC1 cluster in the ‘IL15’ scRNA-seq dataset.

**Supplementary Table 8 (related to Fig. 5):** Marker genes for the clusters defined in Figure 5 in the ‘All Cells’ scRNA-seq dataset.

## Acknowledgments

We thank the UC Irvine Genomics Research and Technology (GRT) Hub (M. Oakes, S. Chung, Q. Nguyen, and V. Ciobanu) for their technical assistance with single-cell library sequencing, and the UC Irvine Sue and Bill Gross Stem Cell Research Center Flow Core (V. Scarfone and P. Nguyen) for their help and assistance with flow cytometry and fluorescence-activated cell sorting. We are also grateful to Angela Lincy Prem Antony Samy and Tatyana Lev for their support with this study.

## Author contributions

Conceptualization: TPM, PN, KK, TEO, AM, JY, DAL Methodology: TPM, PN, LT, EZh, AJL, IA

Investigation: TPM, PN, EZa, LT, EZh, NW, AJL, IA, JIR, HS, IH, SAG, DM, MHL, SM

Visualization: TPM, PN, EZh, NW

Funding acquisition: TPM, PN, KK, JY, DAL

Project administration: TPM, PN, DAL Supervision: DAL

Writing – original draft: TPM, PN, DAL

## Data and code availability

All data reported in this paper will be shared with the lead contact upon request.

---

**Figure S1.**
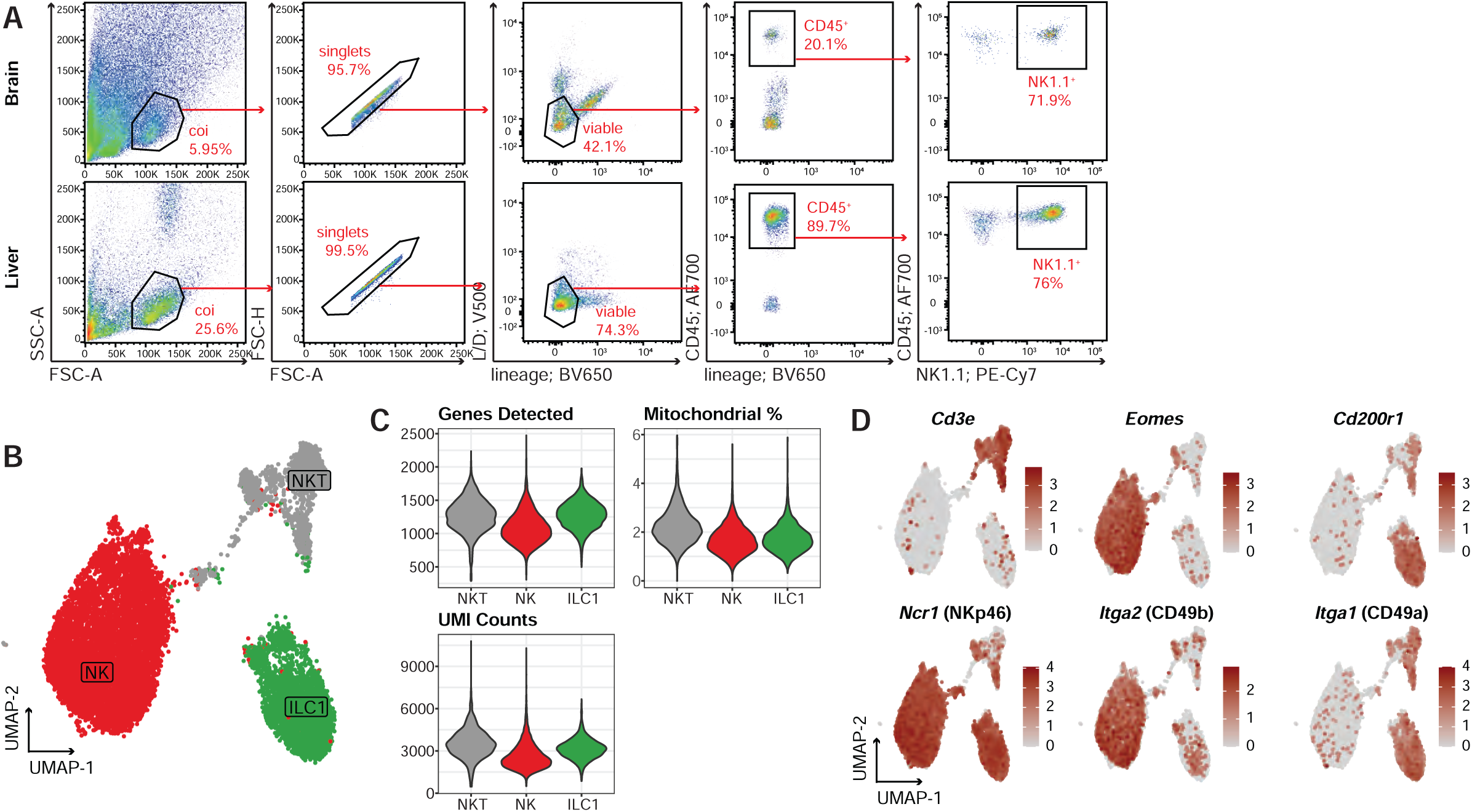
Generating a type 1 ILC-enriched scRNA-seq dataset of healthy and metastasis-burdened livers and brains. Related to Figure 1. **(A)** Flow cytometry gating strategy to sort NK1.1^+^ cells in murine brain (top panel) and liver (bottom panel). After tissue digestion, we enriched for type 1 ILCs by magnetic cell separation and performed lineage depletion (removing CD19^+^, Gr-1^+^, Ter119^+^, EpCAM^+^, CD4^+^, and CD8^+^ cells). Then, we stained the lineage-negative cells for fluorescence-activated cell sorting (FACS). Finally, we sorted viable, CD45^+^NK1.1^+^ cells for scRNA-seq. **(B)** UMAP visualization of scRNA-seq profiles from sorted NK1.1^+^ cells isolated from the brain and liver of PBS-injected (healthy) or tumor-burdened BL/6 mice, including an NK T-cell (NKT) cluster. Data from the TimeCourse dataset. **(C)** Violin plots showing the genes detected, percentage mitochondrial genes, and unique molecular identifier (UMI) counts from NKTs, NKs, and ILC1s. Data from the TimeCourse dataset. **(D)** Feature plots showing expression of *Cd3e*, *Ncr1* (NKp46), *Eomes, Itga2* (CD49b), *Cd200r1, and Itga1* (CD49a). Data from the TimeCourse dataset.

**Figure S2.**
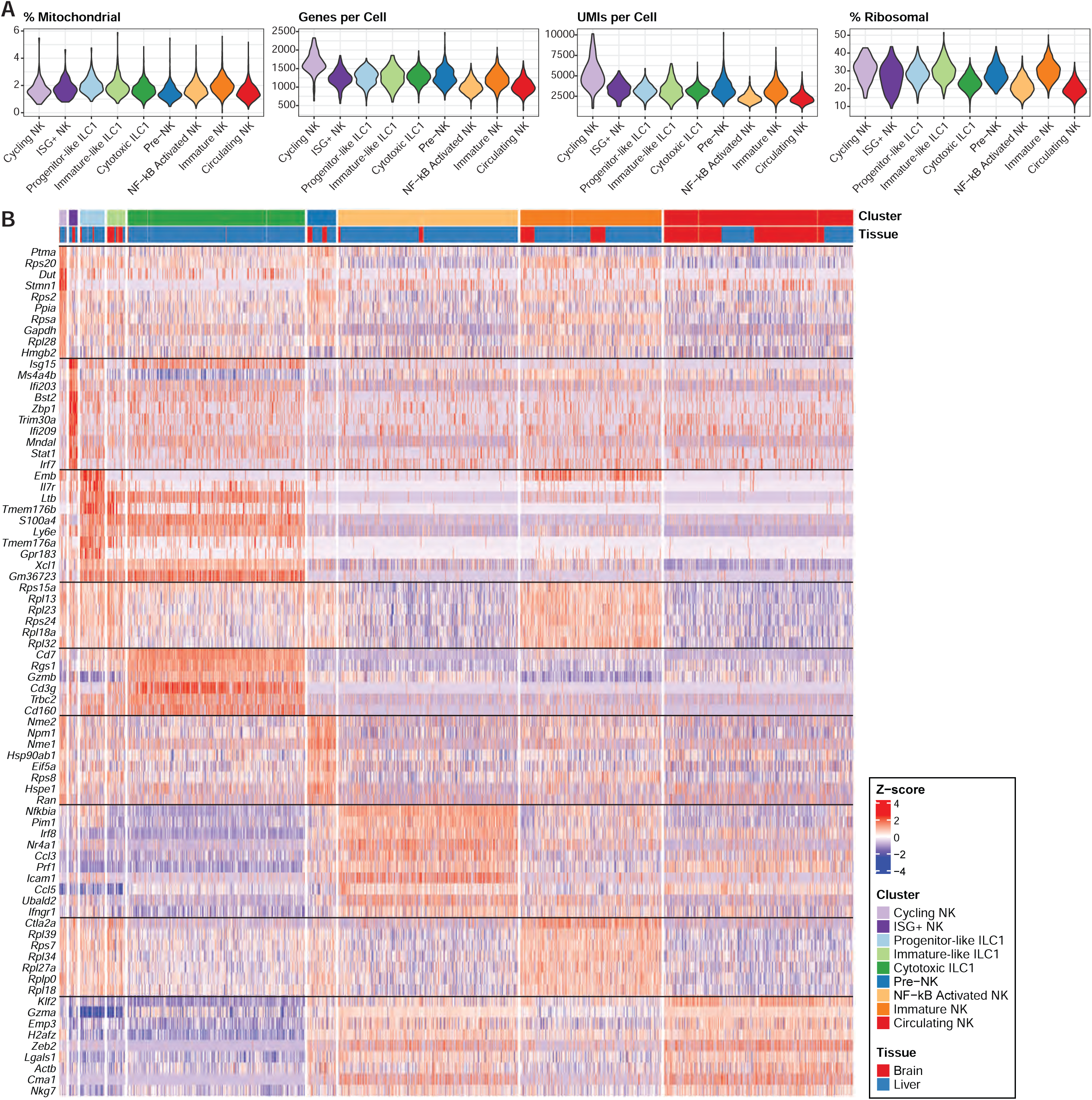
Characterization of the type 1 ILC-enriched scRNA-seq dataset. Related to Figure 1. **(A)** Violin plots showing the percentage of mitochondrial genes, genes per cell, unique molecular identifier (UMI) counts, and percentage of ribosomal genes for the defined NK and ILC1 clusters in Figure 1. **(B)** Heatmap showing the top 10 genes for each of the defined NK and ILC1 clusters in Figure 1. The bars on the top indicate cluster identity and tissue. Values represent Z-scored gene expression.

**Figure S3.**
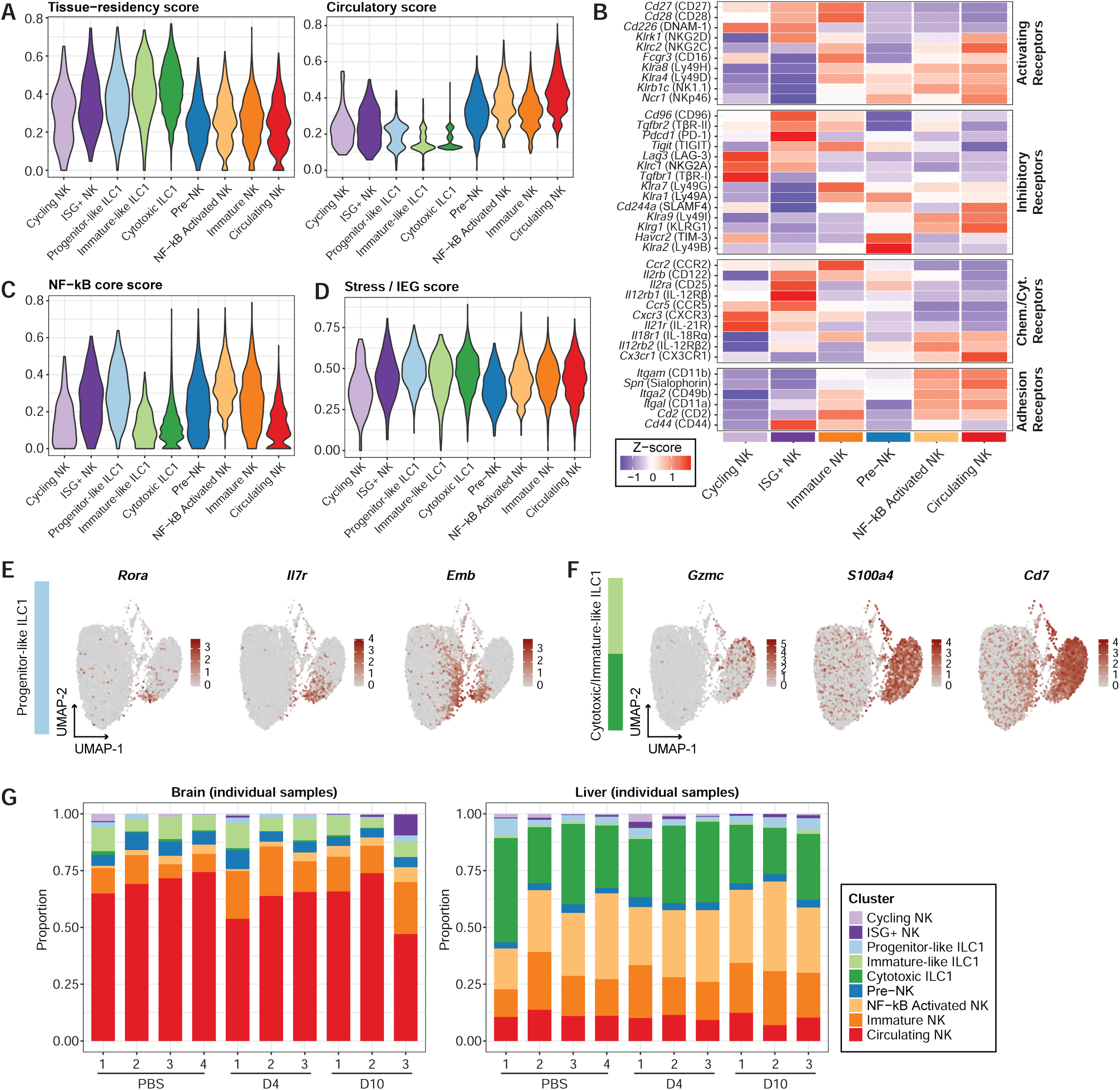
Characterization of the type 1 ILCs identified in the scRNA-seq dataset. Related to Figure 1. **(A)** Violin plots showing the module scores for the tissue-residency and circulatory gene programs in each of the defined NK and ILC1 clusters in Figure 1. **(B)** Heatmap of Z-scored NK cell genes in each of the defined NK cell clusters in Figure 1. We grouped the genes by biological function: activating receptors, inhibitory receptors, chemokine/cytokine receptors, and adhesion receptors. Gene names are italicized; translated protein names are in brackets. **(C-D)** Violin plots showing the module scores for the NF-κB and Stress/IEG gene programs in each of the defined NK and ILC1 clusters in Figure 1. **(E-F)** Feature plots for genes expressed in progenitor-like ILC1s (*Rora*, *Il7r*, *Emb*; E) or cytotoxic/immature-like ILC1s (*Gzmc*, *S100a4*, *Cd7*; F). **(G)** Bar plots showing the cluster distribution in brains (left panel) and livers (right panel) of individual mice, either injected with PBS (Control group) or 4 and 10 days post intracardiac injection of EO771 cells (D4, D10).

**Figure S4.**
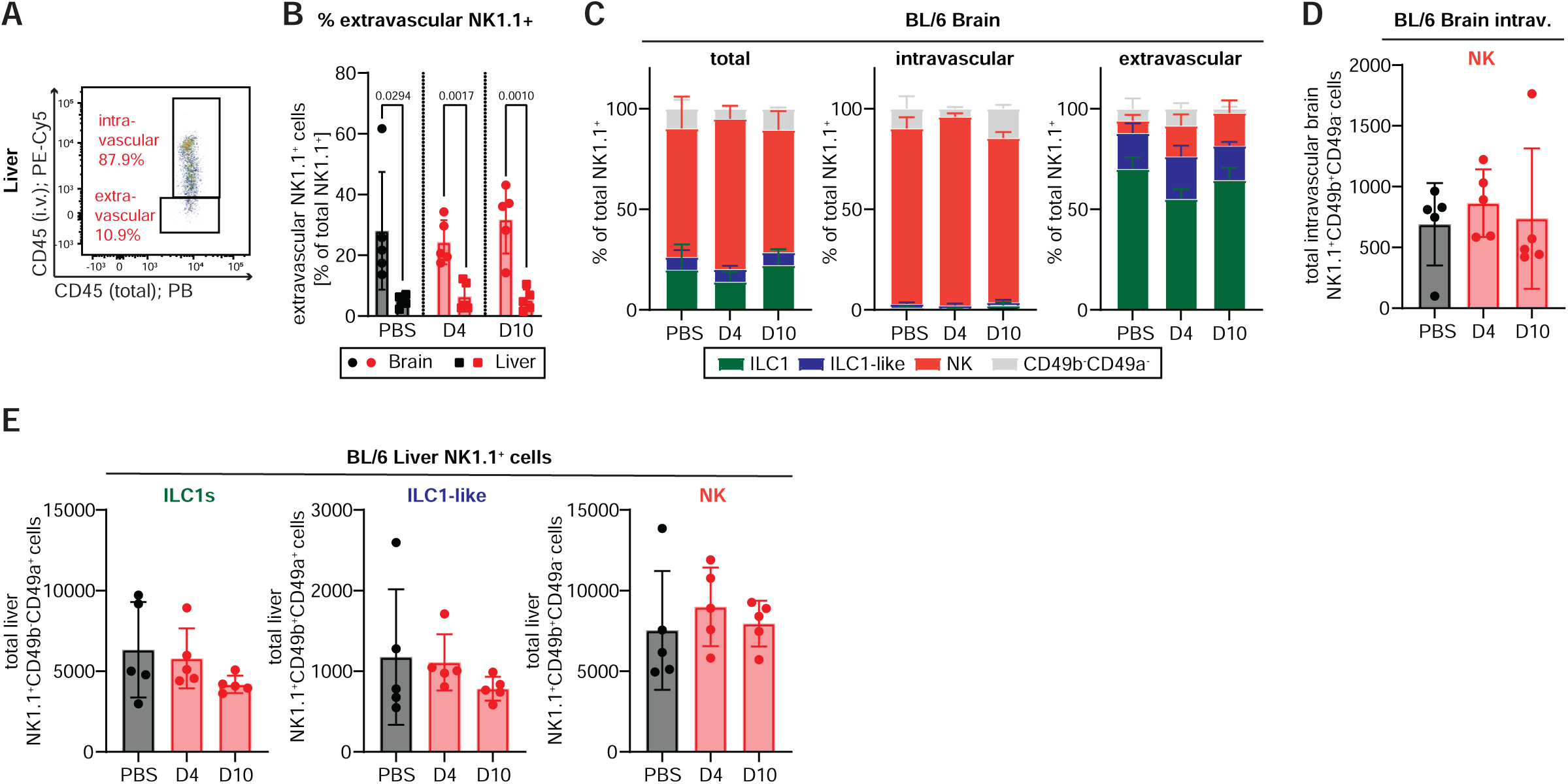
NK cells fail to infiltrate BCBM. Related to Figure 2. **(A)** Flow cytometry gating strategy to identify intravascular (CD45 i.v.^+^/CD45 total^+^) and extravascular (CD45 i.v.^−^/CD45 total^+^) NK1.1^+^ cells in murine liver. Cells were pregated on viable, lineage^-^ (lin^-^; CD19^-^Gr-1^-^Ter119^-^EpCAM^-^CD4^-^CD8^-^), CD45^+^ cells. **(B)** Frequency of extravascular NK1.1^+^ cells (CD45 i.v.^−^/CD45 total^+^) within total NK1.1^+^ cells in liver and brain from PBS-injected controls or metastatic mice at D4 or D10 after intracardiac EO771 eGfly injection. Each dot represents one mouse (n=5 per group and organ). **(C)** Frequencies of ILC1s, ILC1-like cells, and NKs within the total brain NK1.1^+^ cells, the intravascular brain NK1.1^+^ cells, or the extravascular brain NK1.1^+^ cells. **(D)** Absolute counts of intravascular NKs in brains of PBS-(black) or EO771 eGfly-injected BL/6 mice (red; n=5 per group and organ). **(E)** Absolute counts of total ILC1s, ILC1-like cells, and NKs in livers of PBS-(black) or EO771 eGfly-injected BL/6 mice (red; n=5 per group and organ). Data are represented as mean ± SD. Statistics: two-tailed Student’s t-test (B). Exact p-values are shown for statistically significant comparisons.

**Figure S5.**
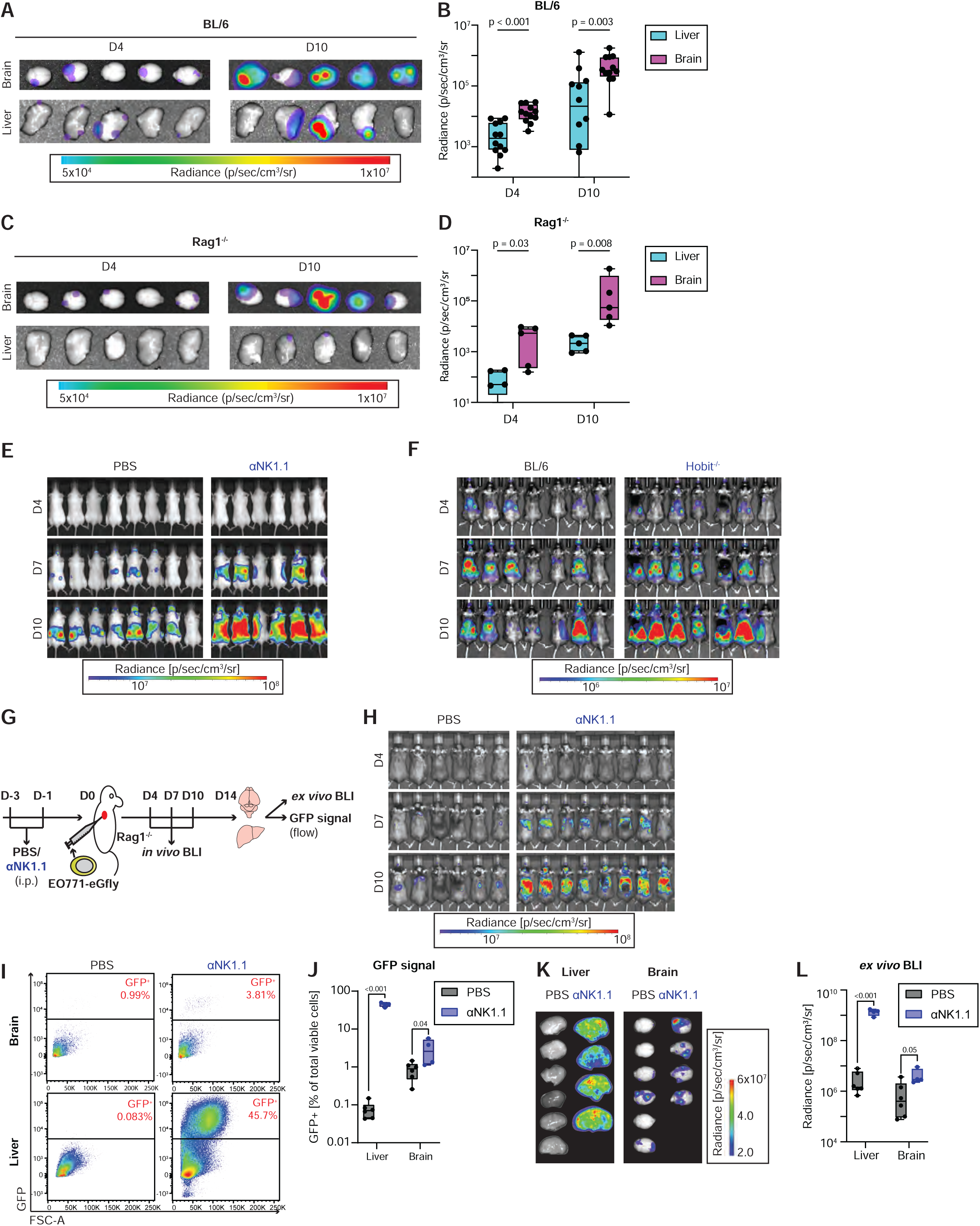
Group 1 ILCs fail to control brain metastasis. Related to Figure 3. **(A-D)** *Ex vivo* tumor burden in brains and livers of BL/6 (A-B) and Rag1^-/-^ (C-D) mice 4 (n=12 for BL/6, n=5 for Rag1^-/-^) and 10 (n=10 for BL/6, n=5 for Rag1^-/-^) days post intracardiac injection of EO711 tumor cells, measured by bioluminescence imaging (BLI). (A) and (C) are the representative images of the metastasis signal in the respective organs; (B) and (D) show the quantification of the metastasis burden. Each dot represents the signal in an individual organ. **(E)** *In vivo* BLI of control (PBS; n=8) and NK1.1-depleted (αNK1.1; n=6) mice at days 4, 7, and 10 after intracardiac EO771 eGfly injection. **(F)** *In vivo* BLI of BL/6 (n=7) and Hobit^-/-^ (n=7) mice at days 4, 7, and 10 after intracardiac EO771 eGfly injection. **(G)** Experimental schematic for systemic NK1.1^+^ cell depletion. Rag1^-/-^ mice received anti-NK1.1 depleting antibody (αNK1.1) 3 and 1 days prior to intracardiac injection of EO771 eGFP/luciferase (eGfly) cells; control mice received PBS instead of αNK1.1. Tumor burden was monitored by bioluminescence imaging (BLI) and flow cytometry, and NK1.1^+^ cell abundance was assessed at the experimental endpoint. **(H)** *In vivo* BLI of control (PBS; n=6) and NK1.1-depleted (αNK1.1; n=8) mice at days 4, 7, and 10 after intracardiac EO771 eGfly injection. **(I-J)** Flow cytometry gating strategy (I) and quantification (J) of GFP^+^ EO771 eGfly tumor cells in digested brains and livers from control (PBS; n=6) and NK1.1-depleted (αNK1.1; n=4) Rag1^-/-^ mice. Cells were pre-gated on viable cells. **(K-L)** *Ex vivo* BLI of brains and livers harvested on day 14 post-injection (K) and corresponding quantification of total bioluminescence signal (L) from control (PBS; n=6) and NK1.1-depleted (αNK1.1; n=4) mice. Data are represented as mean ± SD. Statistics: two-tailed Student’s t-test (B, D, J, L). Exact p-values are shown for statistically significant comparisons.

**Figure S6.**
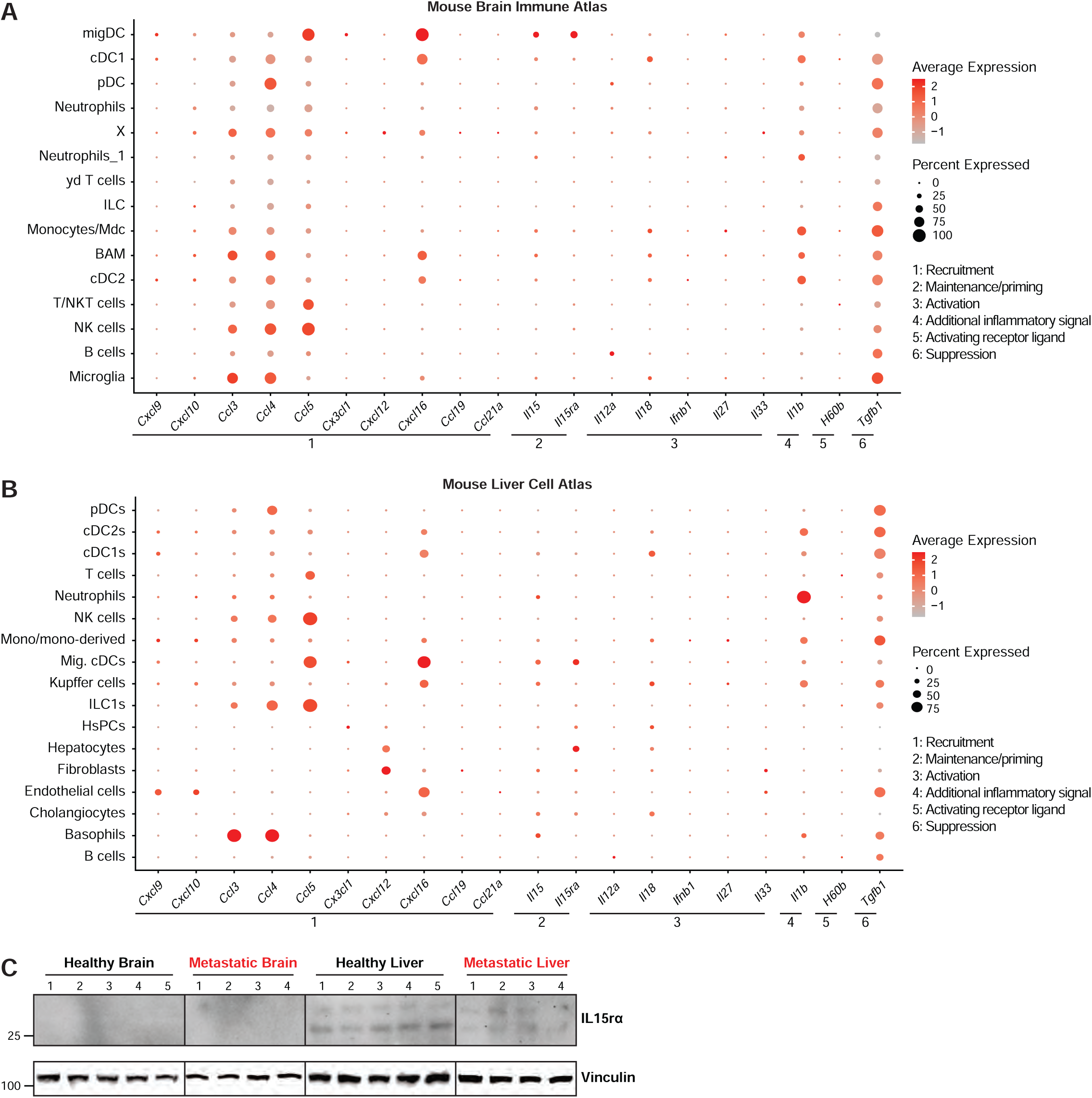
Limited IL15/IL-15Rα expression in the CNS. Related to Figure 4. **(A-B)** Dot plots showing the expression of group 1 ILC recruitment, maintenance/priming, activation, inflammatory signal, activating receptor ligand, and suppression genes in the murine brain and liver. The dot plots have been generated using the Mouse Brain Immune Atlas (GSE128855; A) or the Mouse Liver Cell Atlas (GSE192742; B). HsPCs: hematopoietic stem and progenitor cells; Mig. cDCs: migratory conventional dendritic cells; cDCs: conventional dendritic cells; pDCs: plasmacytoid dendritic cells; migDC: migratory dendritic cells; BAM: border-associated macrophages. **(C)** Western blot showing the expression of IL-15Rɑ and Vinculin in the brain and livers of healthy BL/6 mice or BL/6 animals 9 days post intracardiac injection of EO711-eGFly cells.

**Figure S7.**
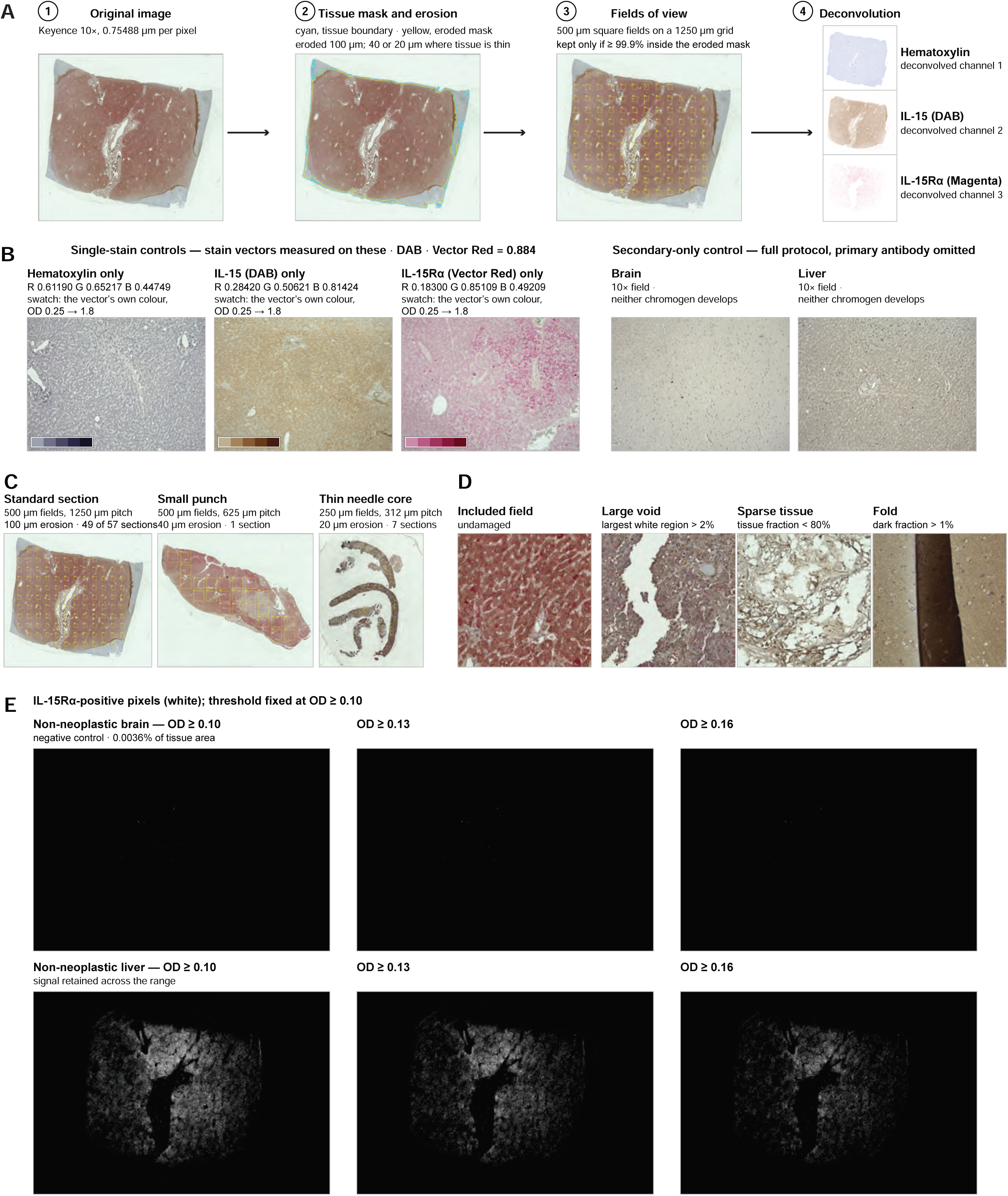
Validation and workflow for quantitative duplex immunohistochemistry analysis of IL-15 and IL-15Rα expression. Related to Figure 4. **(A)** Overview of the image-analysis workflow used to quantify IL-15 and IL-15Rα staining in human brain and liver tissue sections. We acquired whole-slide images at 10× magnification, then used automated tissue segmentation and eroded tissue boundaries to minimize edge artifacts. We then sampled standardized square fields of view (FOVs) across the tissue and retained them only when ≥99.9% of the field fell within the eroded tissue mask. We then deconvoluted images into hematoxylin, IL-15/DAB, and IL-15Rα/magenta channels for independent quantification. **(B)** Validation of chromogen deconvolution and staining specificity. We used single-stain controls for hematoxylin, IL-15/DAB, and IL-15Rα/magenta to define stain vectors for color deconvolution. Secondary-only controls from brain and liver tissue, processed using the complete staining protocol but omitting primary antibodies, showed no appreciable chromogen development. **(C)** Sampling strategy for sections of different sizes. Standard tissue sections were analyzed using 500-µm FOVs positioned on a 1,250-µm grid with 100-µm tissue-mask erosion. For small tissue sections, grid spacing was reduced to 625 µm, whereas thin needle-core sections were analyzed using 250-µm FOVs on a 312-µm grid with 20-µm erosion. **(D)** Representative examples of FOV quality control. Undamaged tissue fields were included, whereas fields containing excessive void space, tissue tears, or folds were excluded according to predefined image-quality criteria. **(E)** Validation of the optical-density threshold used to define IL-15Rα-positive pixels. Representative non-neoplastic brain and liver sections are shown across increasing optical-density thresholds. We selected an OD threshold of ≥0.10 for subsequent quantification, retaining robust IL-15Rα signal in the liver while producing minimal positive area in non-neoplastic brain. White pixels indicate regions classified as IL-15Rα positive.

**Figure S8.**
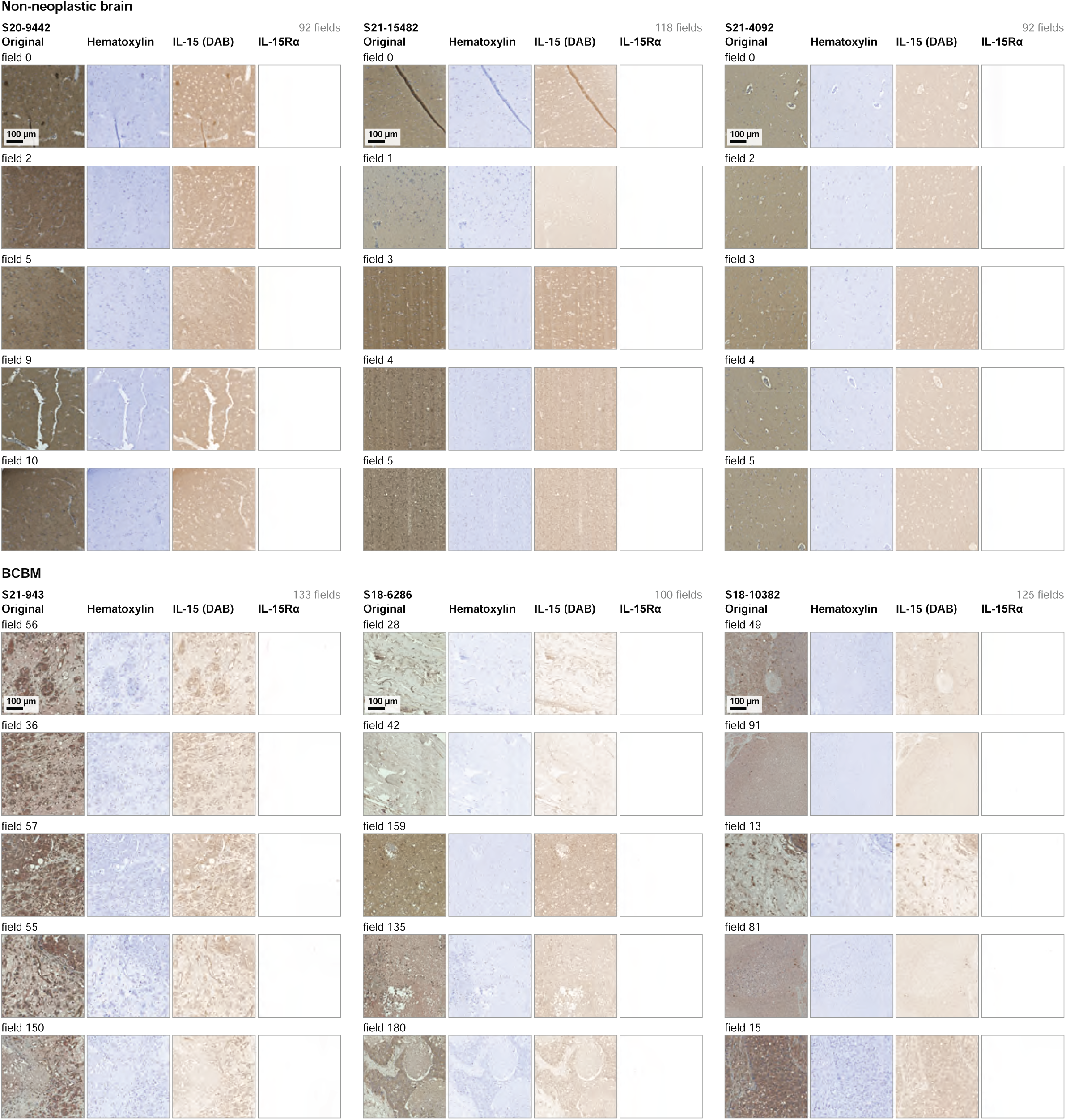
Limited IL15/IL-15Rα expression in the CNS. Related to Figure 4. Micrographs show representative FOVs with complete staining (original) or after deconvolution into the hematoxylin, IL-15 (DAB), and IL-15Rɑ (magenta) channels for 3 non-neoplastic brain sections (S20-9442, S21-15482, S21-4092), and 3 BCBM sections (S21-943, S18-6286, S18-10382). We show five representative FOVs per section. The percentage of IL-15Rɑ-positive area is shown for each FOV, as well as the median for all FOVs per section. Scale bar: 100 µm.

**Figure S9.**
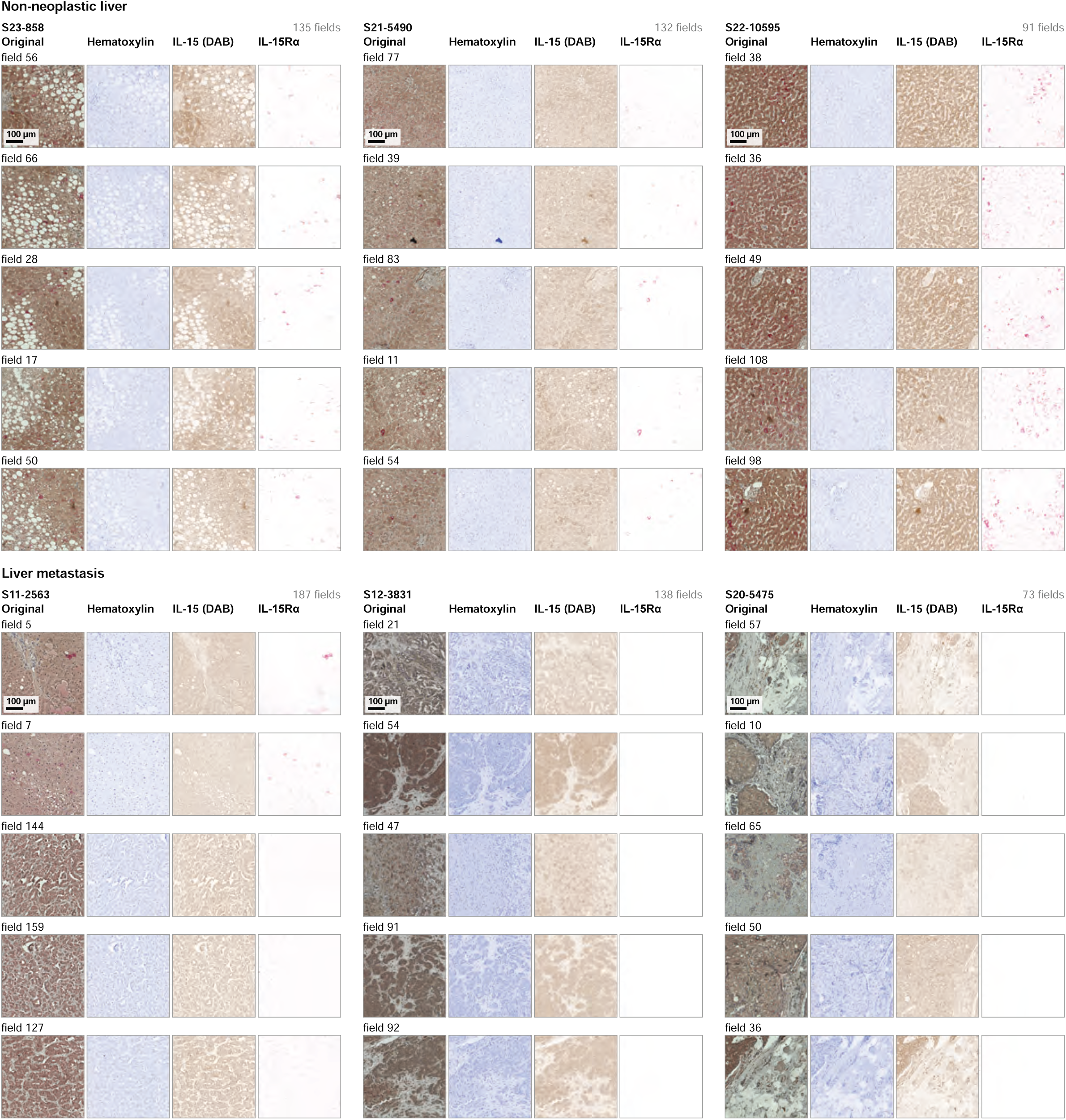
Limited IL15/IL-15Rα expression in the CNS. Related to Figure 4. Micrographs show representative FOVs with complete staining (original) or after deconvolution into the hematoxylin, IL-15 (DAB), and IL-15Rɑ (magenta) channels for 3 non-neoplastic liver sections (S23-858, S21-5490, S22-10595), and 3 liver metastasis sections (S11-2563, S12-3831, S20-5475). We show five representative FOVs per section. The percentage of IL-15Rɑ-positive area is shown for each FOV, as well as the median for all FOVs per section. Scale bar: 100 µm.

**Figure S10.**
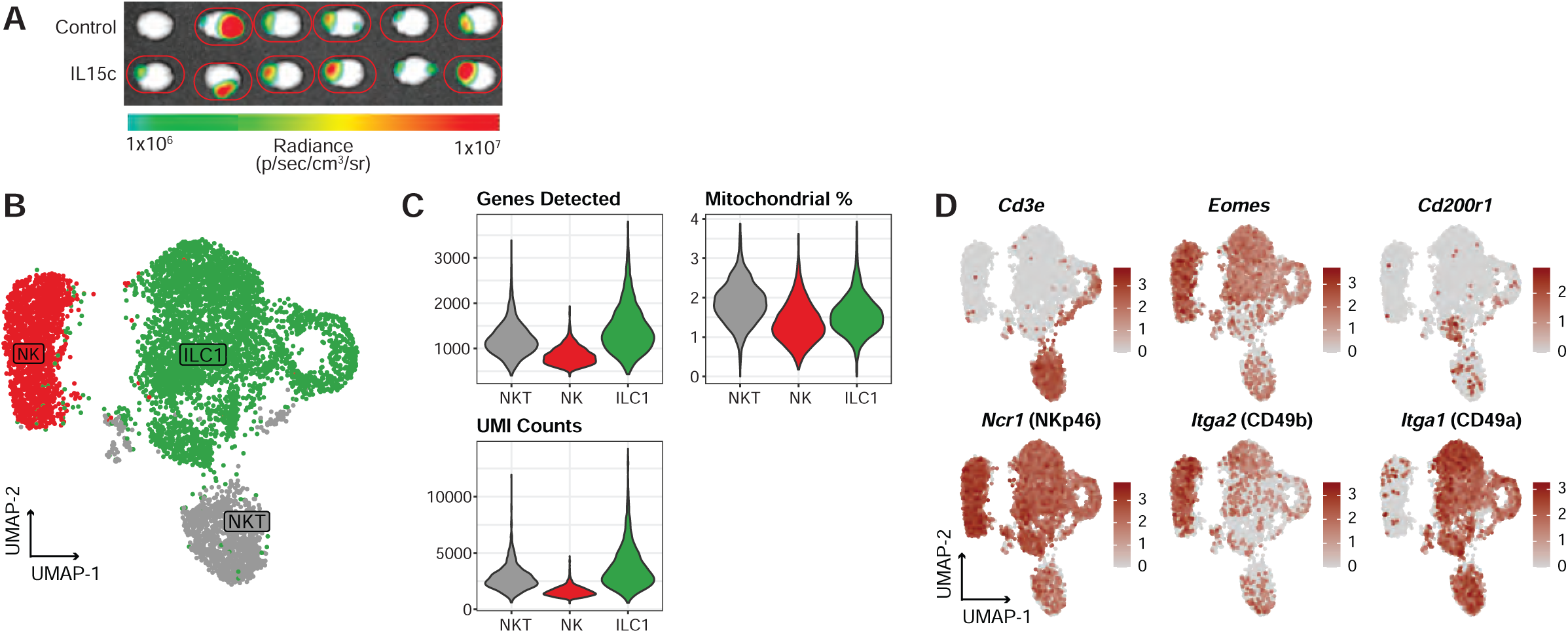
Generating a type 1 ILC-enriched scRNA-seq dataset of metastatic brains from PBS- and IL-15c-treated mice. Related to Figure 5. **(A)** *Ex vivo* tumor burden in metastatic brains of BL/6 mice that were treated with PBS (control) or IL-15 complex (IL-15c). Red circles indicate the brains selected for sequencing. **(B)** UMAP visualization of scRNA-seq profiles from sorted NK1.1^+^ cells isolated from the brain of PBS- or IL-15c-treated metastasis-burdened BL/6 mice, including an NK T-cell (NKT) cluster. **(C)** Violin plots showing the genes detected, percentage mitochondrial genes, and unique molecular identifier (UMI) counts from NKTs, NKs, and ILC1s. **(D)** Feature plots showing expression of *Cd3e*, *Ncr1* (NKp46), *Eomes, Itga2* (CD49b), *Cd200r1, and Itga1* (CD49a).

**Figure S11.**
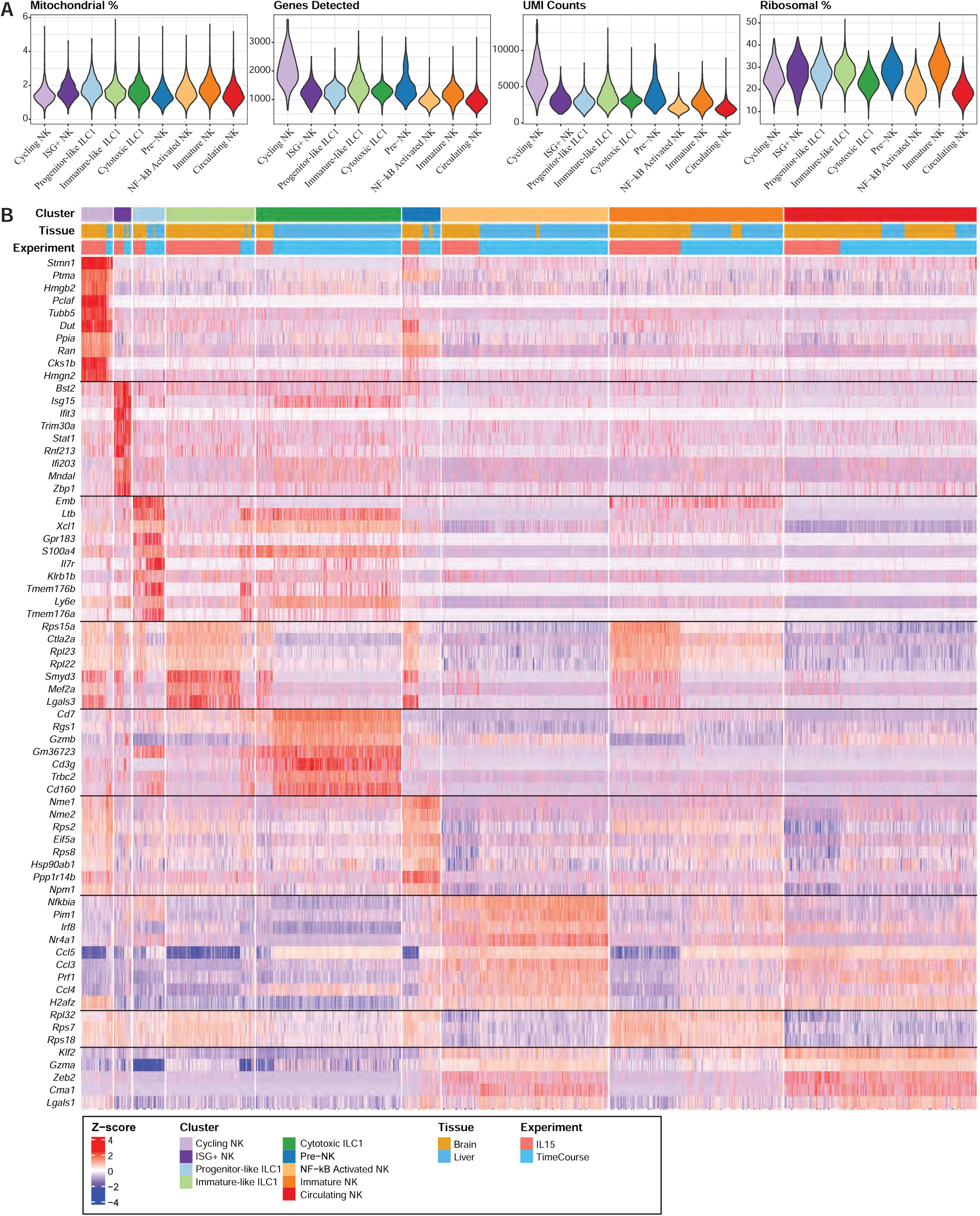
Characterization of the type 1 ILC-enriched scRNA-seq dataset. Related to Figure 5. **(A)** Violin plots showing the percentage of mitochondrial genes, genes per cell, unique molecular identifier (UMI) counts, and percentage of ribosomal genes for the defined NK and ILC1 clusters in Figure 5. **(B)** Heatmap showing the top 10 genes for each of the defined NK and ILC1 clusters in Figure 5. The bars on the top indicate cluster identity, tissue, and experiment. The experiment bar is split into ‘TimeCourse’, originating from Fig. S1, and ‘IL15’, originating from Fig. S10. Values represent Z-scored gene expression.

**Figure S12.**
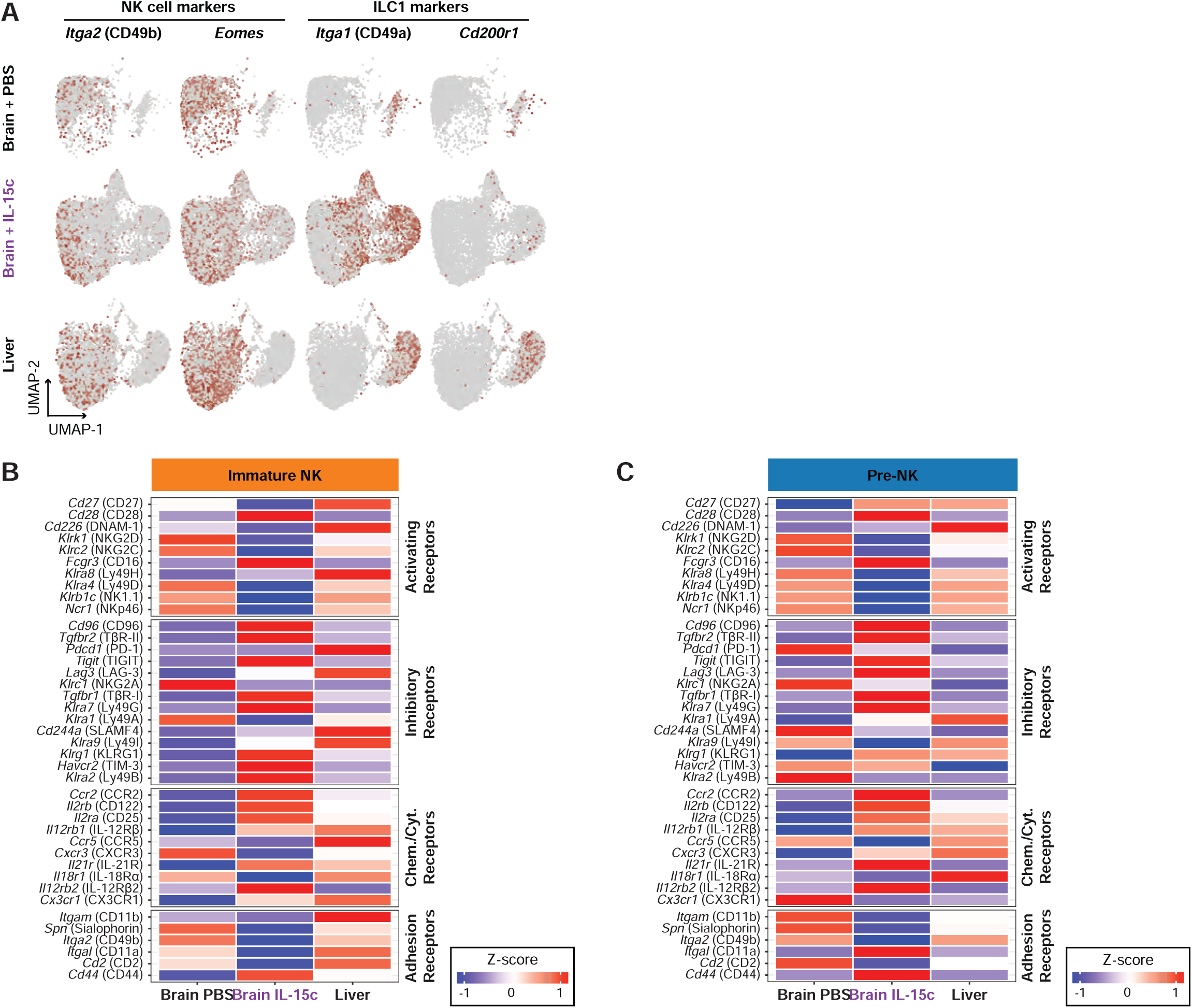
Characterization of the type 1 ILC-enriched scRNA-seq dataset. Related to Figure 5. **(A)** Feature plots depicting expression of *Itga2* (CD49b), *Eomes*, *Itga1* (CD49a), and *Cd200r1* across NK1.1^+^ cells, split by tissue and treatment. **(B-C)** Heatmap of Z-scored NK cell genes in the “Immature NK” (B) and “Pre-NK” clusters, split by the Brain PBS, Brain IL-15c, and Liver groups. We grouped the genes by biological function: activating receptors, inhibitory receptors, chemokine/cytokine receptors, and adhesion receptors. Gene names are italicized; translated protein names are in brackets.

**Figure S13.**
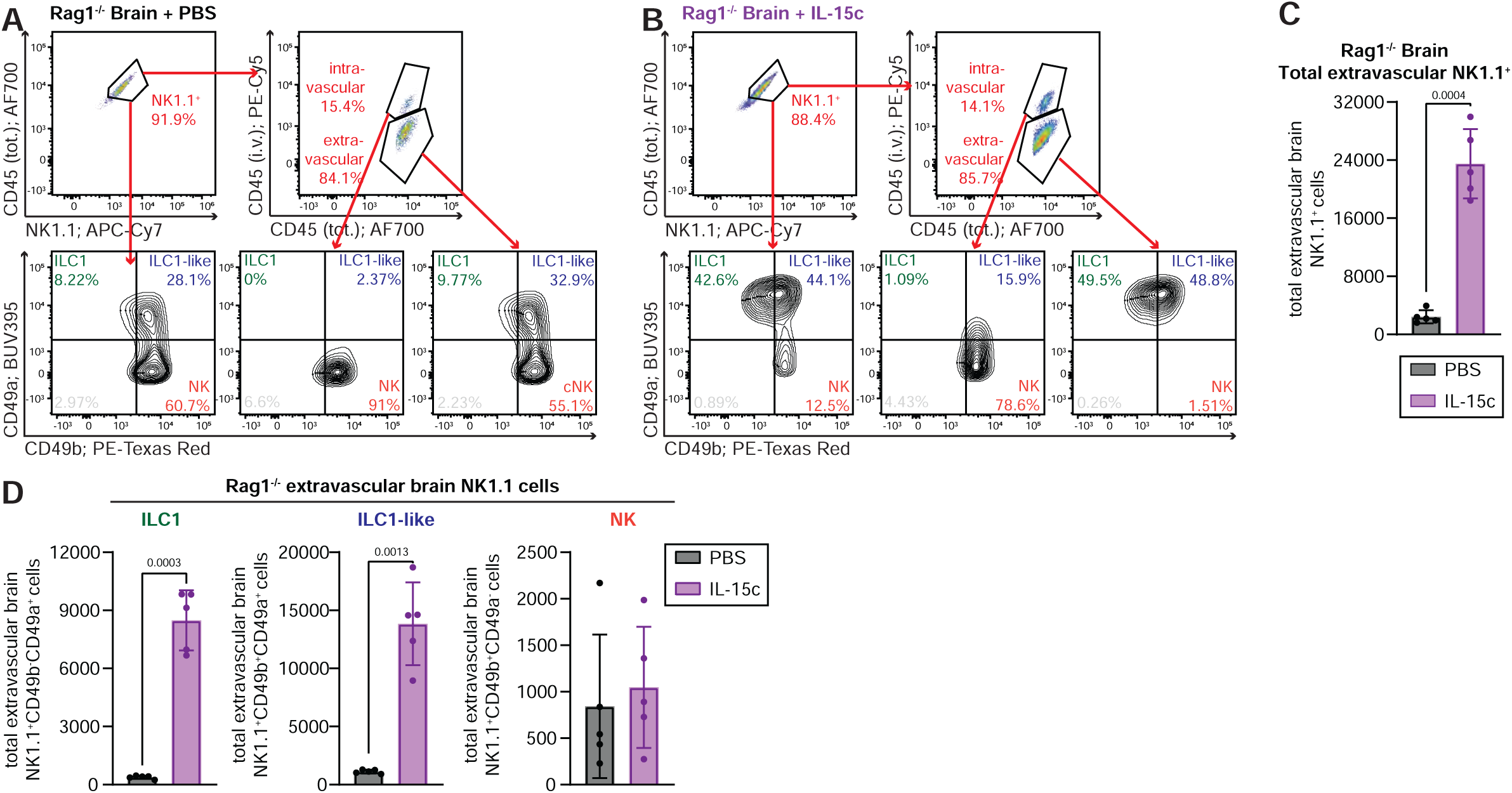
IL-15/IL15Ra rescues group 1 ILC diversity and expands effector populations in BCBM. Related to Figure 5. **(A-B)** Flow cytometry gating strategy to identify intra- and extravascular brain CD49a^+^ ILC1s, CD49a^+^CD49b^+^ ILC1-like cells, and CD49b^+^ NKs in EO771 eGfly–injected Rag1^-/-^ mice treated with PBS (A) or IL-15c (B). Mice received three doses of PBS or IL-15c (D0, D1, and D4) and were sacrificed on day 7. **(C-D)** Absolute numbers of total extravascular NK1.1^+^ (C), ILC1s, ILC1-like cells, and NKs (D) in brains of Rag1^-/-^ mice that were intracranially injected with EO771 eGfly cells, in combination with PBS (n=5) or IL-15c (n=5). Mice received three doses of PBS or IL-15c (D0, D1, and D4) and were sacrificed on day 7. Data are represented as mean ± SD. Statistics: two-tailed Student’s t-test (C, D). Exact p-values are shown for statistically significant comparisons.

**Figure S14.**
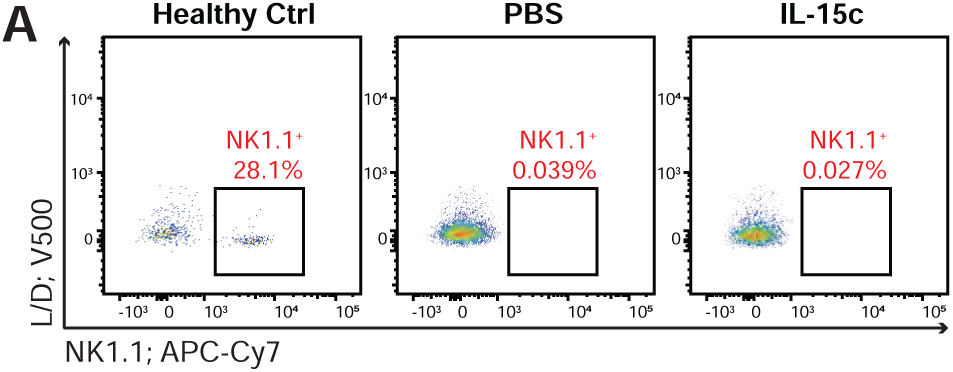
Group 1 ILCs are critical to limit brain metastasis upon IL-15c treatment. Related to Figure 6. **(A)** Flow cytometry gating strategy showing the abundance of NK1.1^+^ cells in the brain of a healthy control BL/6 mouse (left) and NK1.1-depleted BL/6 mice (middle and right panels). The mouse in the middle (PBS) and right (IL-15c) panels received an intracranial injection of EO771 cells (D0) and were then treated with PBS (middle panel) or IL-15 complex (IL-15c, right panel) on days 0, 1, and 4. We depleted NK cells with an anti-NK1.1 antibody 3 and 1 day before injecting EO771 cells. Mice were sacrificed on day 7.

**Figure S15.**
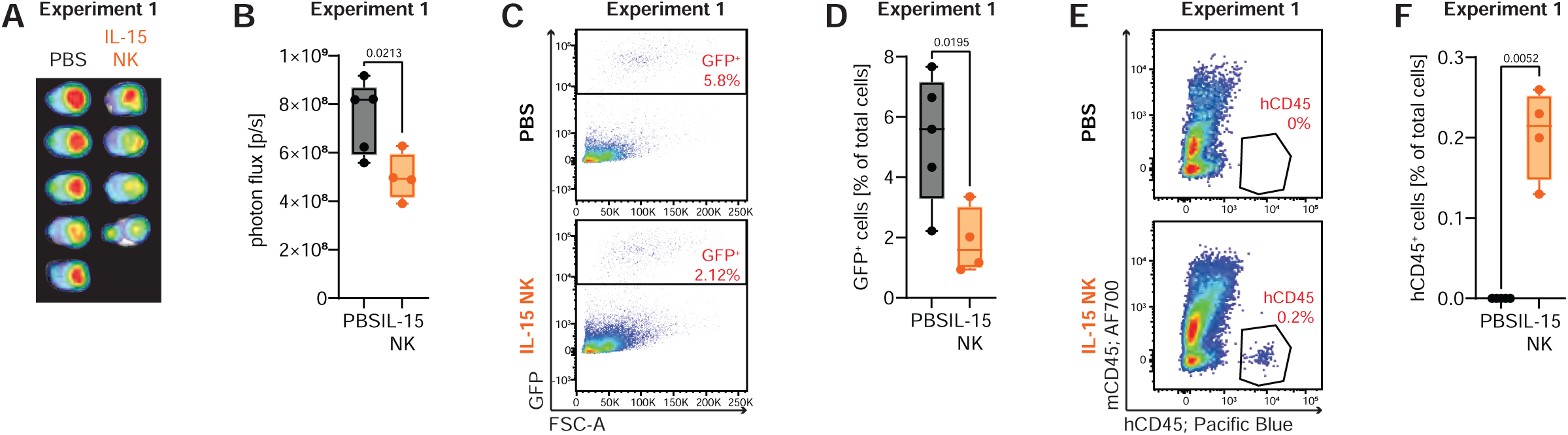
iPSC-derived NK cell therapy improves control of brain metastasis. Related to Figure 7. **(A-B)** *Ex vivo* BLI of brains harvested on day 15 post-implantation (A) and corresponding quantification of total bioluminescence signal (B) from PBS-treated controls and iPSC-derived NK cell–treated mice. **(C-D)** Flow cytometry gating strategy (C) and quantification (D) of GFP^+^ MDA-MB-231 eGfly tumor cells in digested brains from PBS-treated controls and iPSC-derived NK cell–treated NSG mice. Cells were pre-gated on viable cells. **(E-F)** Flow cytometry gating strategy (E) and quantification (F) of human NK cells (mCD45^−^hCD45^+^) in brains from PBS-treated and iPSC-derived NK cell–treated NSG mice. Cells were pre-gated on viable cells; data are shown as the frequency of hCD45^+^ cells among all viable cells. Data are represented as mean ± SD. Statistics: two-tailed Student’s t-test (B, D, F). Exact p-values are shown for statistically significant comparisons.

